# Toward mastering the cell language by learning to generate

**DOI:** 10.1101/2024.01.25.577152

**Authors:** Yixin Chen, Haiyang Bian, Lei Wei, Jinmeng Jia, Xiaomin Dong, Yuzhuo Li, Yubo Zhao, Xinze Wu, Chen Li, Erpai Luo, Chuxi Xiao, Minsheng Hao, Xuegong Zhang

## Abstract

Cells can be viewed as complex stories written by coordinated expression of genes. The success of AI large language models (LLMs) in mastering the human language inspired us to develop a large AI model scMulan with 368 million parameters to generate cell transcriptomics with designated attributes by learning the cell language. We defined a unified c-sentence to incorporate cell transcriptomics and meta-attributes, and pre-trained scMulan on the equivalence of 100 million human cells. Experiments showed that scMulan can generate designated pseudo transcriptomics, predict missing attributes of cells, reconstruct unobserved cells along functional gradients, and can help to identify driving regulators of cell fates. The generated data passed tests of current tools and can reflect the underlying biology.

## Introduction

Understanding the complex mechanism of cells is fundamental in biology and medicine. Cells possess a myriad of attributes, such as the expression levels and patterns of all genes, cell types and states, and other attributes related to their origins. These diverse attributes and their intricate interrelationships can be envisioned as the “cell language”, where each attribute represents a “word” with its “semantics”, and the interactions among them form the “grammar” that defines biological rules. Understanding the mechanism of cells is like deciphering this language, which is crucial for understanding cellular heterogeneity, functions, and behaviors, as well as unraveling disease processes and developing effective therapies.

With the rapid development of new technologies such as single-cell RNA-sequencing (scRNA-seq), scientists are able to obtain snapshots of single-cell transcriptomics of many cells with a variety of other attributes. As genes work as complex systems rather than individuals, deciphering the full gene expression language behind complex cellular phenotypes and subtle variations needs a systematic approach beyond finding cell clusters, trajectories and their marker genes. There even lacks a formal, quantitative way for representing the complex relations of multiple genes and complex cellular attributes, nor an effective way to experimentally validate possible high-order associations or causal relations between genes and cell attributes. With the millions to billions of single cells from all human organs being sequenced for their transcriptomics profiles, the community are accumulating massive “cell language corpora” of virtually all cell types and cell states of the human body in health and diseases. If we treat genes and other attributes of a cell as “tokens” in the language, the size of all available human scRNA-seq data in terms of the total number of tokens is already comparable with the natural language data used to pre-train those large language models such as GPT or Llama^1–4^. This observation suggested the question whether it would be possible to invent large AI models to learn from the large corpora to master the cell language.

Traditional AI studies for natural language understanding were to develop methods for specific types of tasks in semantics and grammar analyses^5,6^. They had achieved big advancement in some specific tasks, but can hardly scale up to comprehensive tasks such as generating human-level comprehensive text contents. LLMs based on the Transformer structure with deep attention mechanism showed that by pre-training the model with sufficiently rich data for self-supervised tasks such as masked word or next word prediction, the models can perform like foundation models that are capable for many challenging downstream tasks which require real “understanding” of the language, such as generating long dialogs, writing essays, or even doing multiple-step reasoning^7–9^. This breakthrough in natural language understanding and in other fields of AI such as multimedia generation suggests the possibility of a new AI paradigm for scientists’ efforts in understanding the biological language of cells, beyond current efforts of probing specific biological questions from single-cell data using specific models.

Current single-cell RNA-seq studied have enabled many important discoveries related with cell types and cell states. Such discoveries are all in the form of qualitative descriptions and graphical illustrations, and each discovery is confined to a very specific biological context, such as the expression patterns of a few genes defining a common or rare cell type that may play a key role in a certain biological process. It is questionable whether or how such discoveries can eventually form the comprehensive systematic understanding of the mechanism behind the cell’s transcriptomic variations and their associated phenotypical variations. Recently, several AI large cellular models (LCMs) have been developed to learn the language of cells using single-cell transcriptomics data, inspired by the success of LLMs on natural languages^10–13^. They are pre-trained on massive amount of unannotated scRNA-seq data in a self-supervised manner, and have preliminarily showcased the potential to be foundation models by performing well on many downstream tasks from cell type clustering, annotation, gene regulation inference to drug-sensitivity and perturbation response prediction. Their success highlighted a possible new approach or new paradigm to decipher the biology of cells: to design large AI models to learn the general language of cells beyond learning to do specific tasks.

Existing LCMs used the “masked language modeling” mode of pre-training^14–19^ that was initiated by the method BERT in LLMs^20^: Genes with their expressions in cells are converted as “tokens” in the cell language, with each cell as one sentence. The tokens are randomly masked and the LCMs are trained to recover the masked tokens. Such self-supervised pre-training makes LCMs be able to learn the mathematical vector embeddings of the genes and their expressions in the contexts of various cells, and thereafter form vector embeddings of cells based on the gene embeddings. Such embeddings of genes and cells are then used for downstream tasks of analyzing the genes and cells, usually with fine-tuning of the LCMs or binging in an extra model based on the embeddings. These works showed the pre-trained models have learned more information from the context of many cells than the vanilla gene expression of each cell. But existing LCMs are still not yet ready for generating full gene expression profiles for cells with designated types and attributes because they were not designed for true generative pre-training tasks.

To master a language, one must be able to write in the language from scratch to form sentences with intended meanings. Similarly, we perceive that making machines to be able to “intentionally” generate the transcriptomics and other essential attributes of cells is crucial for them to master the language of cells. We developed a large single-cell multi-task language model (scMulan) to explore this possibility. We designed a unified cell language framework that incorporates the expression of genes and all other available attributes of cells as a unified cell sentence or “c-sentence”, and designed a pre-training strategy of true generative manner. This design allows the model to be able to perform all tasks as particular instances of the general task of generating unobserved attributes conditional on given or designated attributes. The current version of the model possesses 368 million learnable parameters to facilitate the grasping of the complexity of the cell language. We carefully curated more than 10 million single-cell transcriptomic profiles of human healthy cells with unified annotation and standardized metadata information, and developed a strategy to augment the data by the scale of 10 to make an equivalence of 100 million cells to pre-train the model. Experiments showed that scMulan not only perform standard single-cell analysis tasks such as cell type prediction in purely “zero-shot” manner, but also can generate the transcriptomics of cells of types or states that have not been observed along certain temporal or functional processes or gradients, and can be used to infer critical genes whose expression change can transform cell state and cell fate. In this way, scMulan can be seen as having illustrated the potential for generating live cells from snapshots. The generated cells can pass the test of current tools for single-cell analysis as “Turing-like test”, which implies that the model has “mastered” the cell language to a great extent. This provides a prototype of building virtual tissues and biological processes with generated “digital cells” which can enable future “*in data*” experiments.

## Results

### Overview of scMulan

scMulan is a large-scale pre-trained generative model for the language of single-cell transcriptomics, built on a special Transformer decoder-only architecture with an attention mechanism^21^. We designed a unified form of structured cell sentences, or “c-sentences”, to operationalize the cell language and developed a pre-training strategy to complete partially given c-sentences. The design allows scMulan to handle multiple tasks as specific cases of the general task of generating unobserved attributes conditioned on given or designated attributes within a c-sentence.

A c-sentence is a sequence where each “word” represents a specific attribute of the cell (**Fig. 1A**, Methods). Each word is structured as a 2-tuple, corresponding to an attribute and its associated value (if applicable). This tuple can denote a gene and its expression level, a type of other attributes of the cell (such as annotated cell type, donor age, donor gender, source organ, etc., referred to here as “meta-attributes” for convenience) paired with its value, or an analytical task token accompanied by a placeholder.

**Fig. 1.**
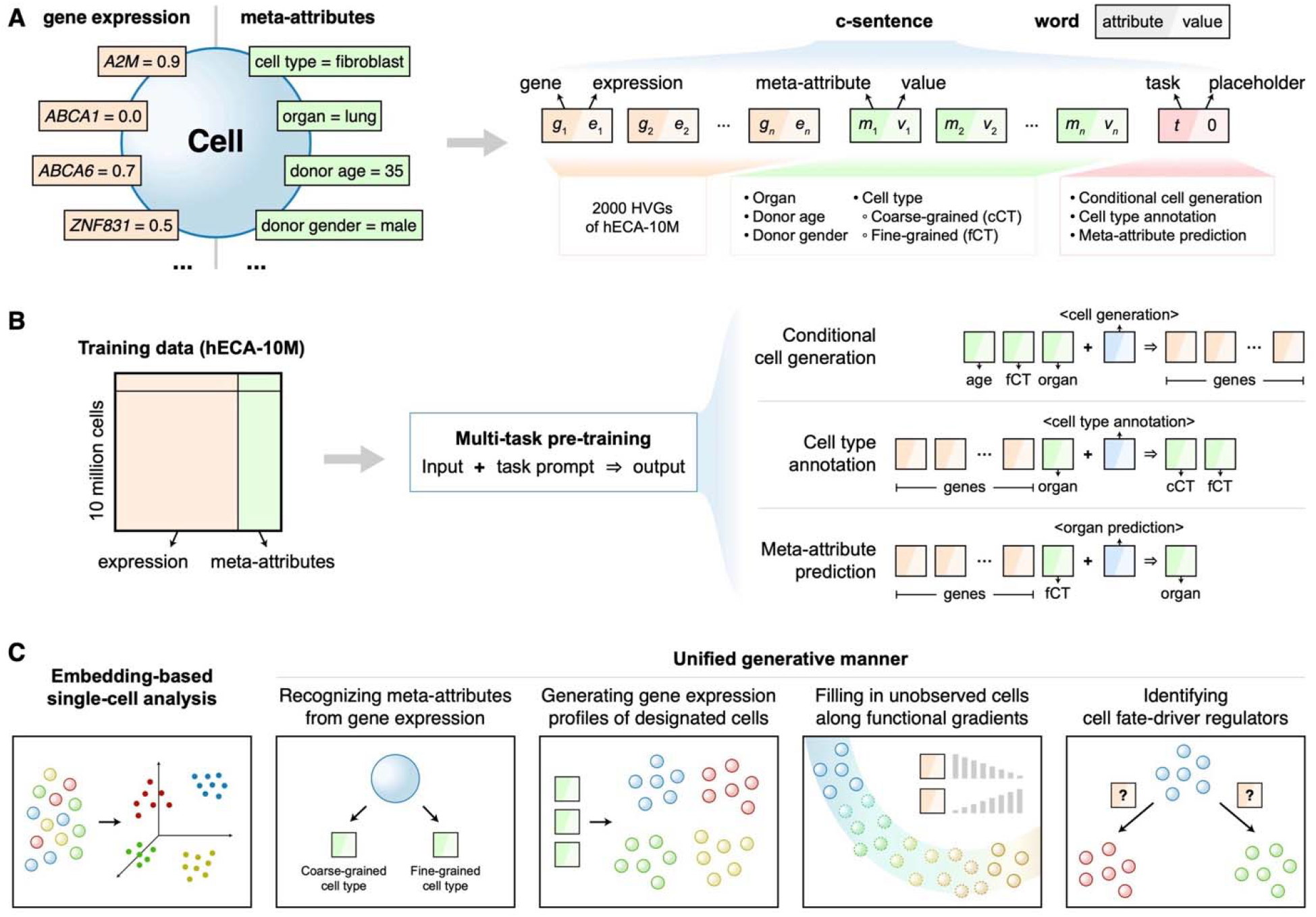
Overview of scMulan. (**A**) The design of c-sentences. Each c-sentence encodes the information of a cell, with each word representing an attribute paired with its value. A word is a 2-tuple which can be a gene and its expression level, a type of meta-attributes paired with its value, or an analytical task token accompanied by a placeholder. (**B**) The workflow of scMulan. scMulan was pre-trained on the hECA-10M dataset with multiple tasks, including conditional cell generation, cell type annotation, and meta-attribute prediction. (**C**) Downstream applications of scMulan. The cell embeddings derived by scMulan can be used for single-cell analysis, and scMulan can perform various downstream tasks in a unified generative manner.

The design of c-sentences enables scMulan to be consistently trained and applied using a unified generative approach. During training, a c-sentence functions similarly to a sentence in natural language processing, where the model can extend a partially provided c-sentence based on the part given. In this way, scMulan can effectively capture the complex dependencies and interactions within cell language through predictive modeling. In practical applications, this framework provides flexibility in generating biological insights. For example, scMulan can generate gene expressions from meta-attributes or *vice versa*, adapting its output to the given input attributes. This flexibility ensures that whether partial gene expressions or meta-attributes are provided, scMulan can complete the missing information by generating a full c-sentence. This unified approach facilitates a smooth transition from model training to real-world applications, ensuring that “insights” gained during pre-training are directly applicable to enhance scMulan’s performance in deployment.

An analytical task token directs scMulan’s output during both pre-training and application. We preset three types of analytical task tokens: “cell type generation”, “organ generation”, and “gene expression generation” in the current version, allowing the model to adjust its output for the corresponding tasks. Using these task tokens, we developed a series of pre-training tasks to help scMulan learn complex relationships among genes and between genes and various meta-attributes. This provides a framework for linking molecular, cellular and phenotypic information. The pre-training tasks include conditional cell generation, hierarchical cell type annotation, and meta-attribute prediction (**Fig. 1B**). Conditional cell generation is a foundational task, which requires scMulan to generate genes and their expression values based on designated meta-attribute conditions, such as organ/tissue name, cell type, donor age, donor gender, sequencing technology, or any combination of these attributes. This task enforces the model to grasp the nuanced relationships between genes and their associated meta-attribute contexts. Hierarchical cell type annotation requires scMulan to generate coarse-grained and fine-grained cell type annotations for each cell. It enforces the model to learn the hierarchical relationship within cell types and their association with gene expressions, which is crucial for understanding cellular homogeneity and heterogeneity. Meta-attribute prediction, exemplified by tasks like predicting the originating organ region of a cell, requires scMulan to recognize meta-attribute-related expression patterns. This task is essential for embedding an understanding of the physical context of cell behavior within the model. This approach can efficiently handle various downstream tasks in a unified generative manner (**Fig. 1C**) and is readily extendable to other meta-attributes, supporting scMulan’s scalability across diverse biological contexts.

We built scMulan upon a Transformer decoder-only architecture with an attention mechanism^22^ (**Fig. S1A**), inspired by causal language modeling in LLMs for natural language processing. In NLP, such models are typically trained to predict the next token (word) based on prior context. A recent model, tGPT, adopted this approach by ordering genes in descending expression level and predicting the next gene based on the preceding ones^23^. However, unlike natural language, gene tokens within a c-sentence do not have a natural sequential order or intrinsic order of importance. Thus, we designed a new strategy and let scMulan to predict the entire set of subsequent tokens based on part of the tokens (Methods). To ensure scMulan’s insensitivity to word order in c-sentences, we randomly shuffled genes within each c-sentence during pre-training. This strategy reinforces the model’s non-sequential learning ability while serving as a data augmentation technique to enlarge the effective size of training data by a factor of 10, enhancing overall model performance.

We utilized the data of the upgraded version of human Ensemble Cell Atlas (hECA)^24^ to train scMulan (**Fig. S1B**). This dataset, named hECA-10M for convenience, includes over 10 million high-quality single-cell transcriptome data from healthy human organs or tissues such as the heart, brain, lung, liver, bone marrow, blood, and thymus. The equivalence size of this dataset is ∼100M cells after the data augmentation. Each cell in the hECA-10M dataset is characterized by 42,117 gene expressions, along with standardized metadata detailing the organ, donor age, donor gender, and cell type. For the sake of saving computational cost, we identified 2,000 highly variable genes (HVGs) in this dataset, and used the HVGs and all metadata as the tokens of c-sentences in the current version. This design enables the inclusion of multiple tasks in the pre-training of scMulan without the need for further supervised labeling, and provides a natural framework for linking molecular, cellular and phenotypic attributes of cells in a unified model.

The current version of scMulan possesses 368 million learnable parameters to facilitate the grasping of the complexity of the cell language. We utilized four A100 80G GPUs to train scMulan on the equivalence of ∼100M cells over a period of 148 hours. Notably, the training efficiency of scMulan significantly surpasses that of previous large cellular models, making it more accessible to labs with limited computing resources. This enhanced efficiency stems largely from the strategic selection of HVGs and the parallel processing capabilities of Transformer decoders. Users can also select their own sets of genes and data of interest, enabling the rapid training of their customized scMulan models. This adaptability ensures that researchers with varying data and resources can leverage the power of large AI models for their specific fields.

### Grasping the multi-granular heterogeneity of cells with scMulan

After the pre-training of scMulan, we evaluated its capability to grasp at multiple granularity the heterogeneous nature of cells encapsulated in the cell language. For each cell, we generated its cell embedding by inputting a c-sentence comprised of the cell’s gene expression profile and the task token “<cell type generation>”, into scMulan, and then extracted the embedding of the last token from the final layer of scMulan’s Transformer. We visualized them using cells from the validation set of hECA-10M (**Fig. S2**, see Methods for details). We can see that the cell embeddings retained critical information about organ types and cell types while displaying minimal batch effects. These results confirmed that scMulan effectively learned the biological variances among cells.

This capacity allows scMulan to handle the data integration task without the need for additional fine-tuning. To evaluate its performance, we compared scMulan with established data integration methods such as BBKNN^25^, Harmony^26^, Seurat v3^27^, FASTMNN^28^, scVI^29^, and scANVI^30^, along with other pre-trained LCMs such as GeneFormer^15^ (without fine-tuning), scGPT^16^ (with and without fine-tuning), and scFoundation^17^ (without fine-tuning). We using two datasets to evaluate the effectiveness in minimizing batch effects while preserving biological variance. The first dataset^31^ included 32,472 lung cells from 16 donors in three datasets (Lung_Luecken_2022), presenting a simpler batch scenario, while the second^32^ was more complex, consisting of 274,346 immune cells from 18 batches, including cells from both healthy donors and COVID-19 patients with varying degrees of severity (COVID-19_ Lotfollahi_2022). These two datasets have not been included in the hECA-10M data. We utilized the benchmarking method scIB^31^ to compare metrics such as the average biological conservation score (AvgBIO) and the running time (Methods).

The integration results are illustrated in **Fig. 2A**. We can see that for the simpler Lung_Luecken_2022 dataset, scMulan’s performance is comparable to BBKNN and scVI, while slightly weaker than scANVI and the fine-tuned scGPT model. For the more complex COVID-19_ Lotfollahi_2022 dataset, scMulan outperforms all other methods. Among the pre-trained models, only scMulan and scFoundation are capable of handling this task in a zero-shot manner (**Fig. 2A**). Only scMulan, scFoundation and the fine-tuned scGPT model successfully integrated COVID-19_ Lotfollahi_2022, with cells of the same type clustering closely together (**Fig. 2B**). Notably, scMulan requires 1/8 the runtime of scFoundation to reach this goal, and scGPT required an additional 10 hours of fine-tuning to achieve this performance (**Fig. 2C**). These findings highlighted scMulan’s exceptional capability for grasping the biological heterogeneity among different cell types despite of technical batches through zero-shot learning.

**Fig. 2.**
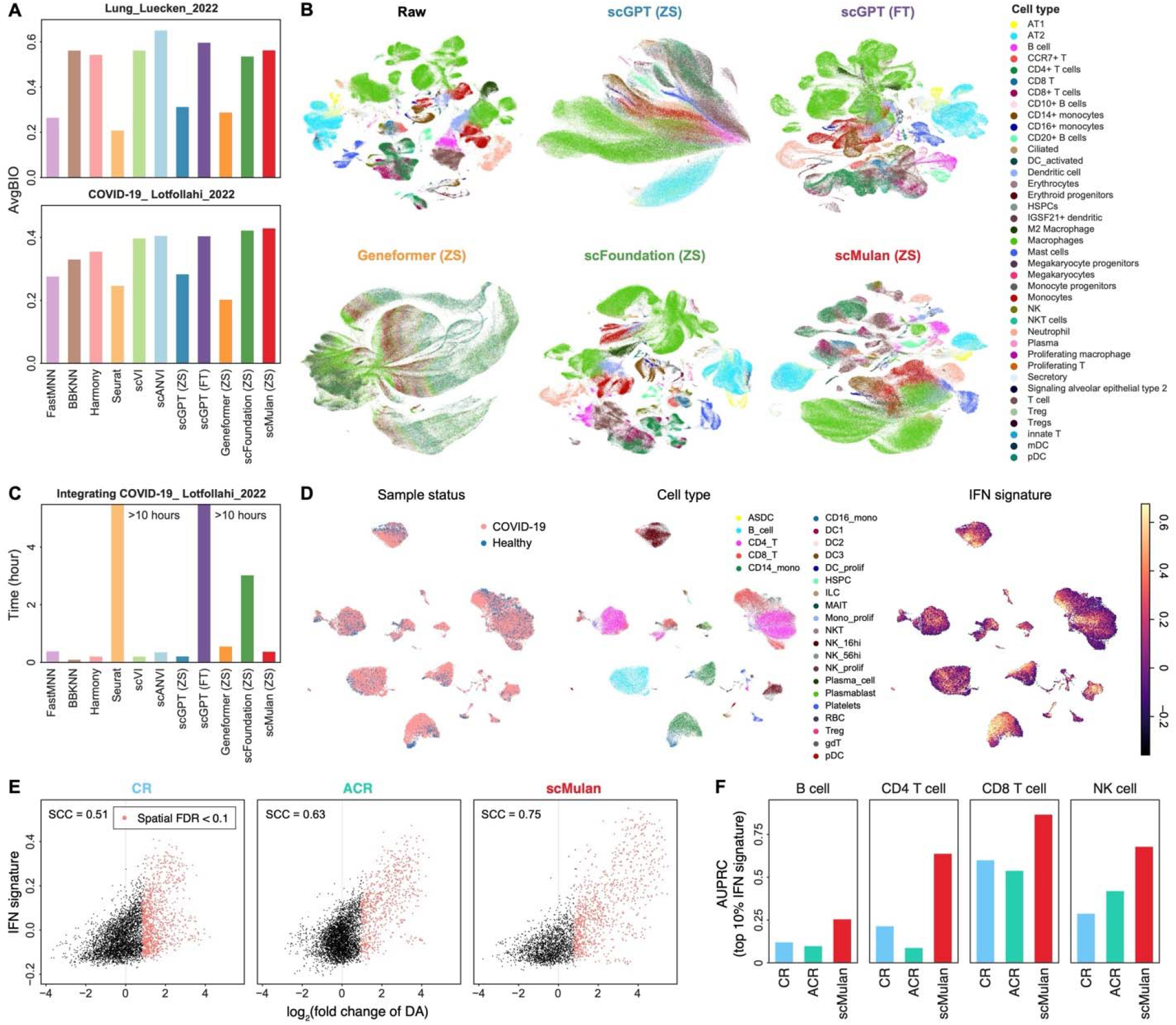
Grasping the multi-granular heterogeneity of cells with scMulan. (**A**) The average biological conservation score (AvgBIO) of different methods in integrating the Lung_Luecken_2022 and the COVID-19_ Lotfollahi_2022 dataset. (**B**) UMAP visualization of COVID-19_ Lotfollahi_2022 with different integration methods. Cells are colored by cell types. (**C**) The running time of different integration methods on COVID-19_ Lotfollahi_2022. (**D**) UMAP visualization of the COVID-19_Dann_2023 dataset with scMulan embeddings. Cells are colored by sample statuses, cell types, and the IFN signature. (**E**) Relationships between DA and the IFN signature for cells in COVID-19_Dann_2023. Cells with neighborhoods significant enriched in COVID-19-associated cells (log_2_(fold change of DA) > 0 and spatial FDR calculated by Milo < 10%) are colored with red. (**F**) AUPRC scores of discriminating cells with high IFN signature by DA for different lymphoid cell types. ZS, zero-shot; FT, fine-tuned; SCC, Spearman’s correlation coefficient.

We then investigated whether scMulan can discern even more nuanced variances across cell states within cell types. This ability is crucial for detecting and analyzing cell state changes during biological processes. We validated this capability with a COVID-19 dataset from a published benchmarking study^33^. This study used a dataset with immune cells from 90 COVID-19 donors and 23 healthy donors (COVID-19_Dann_2023), and they focused on COVID-19-associated cell states which were defined as cells with high IFN signaling profiles. The data were not included in the hECA-10M dataset. The study also defined two paradigms in the joint analysis of single-cell datasets for identifying altered cell states: the control reference (CR) design which integrated the control dataset and the disease dataset for calculating cell embeddings, and an atlas to control reference (ACR) design which mapped the control dataset and the disease dataset onto a cell atlas for calculating cell embeddings. Milo^34^ was employed to detect these states by calculating the neighborhood-level differential abundance (DA) of cells from two groups, and a higher DA score means that the cells in neighbor are mainly from the disease group and more likely to be a COVID-19-associated cell (Methods). We conducted the same experiment on the cell embeddings generated by scMulan and compared its performance against the results provided in the original work.

The UMAP visualization of scMulan embeddings revealed that immune cells from COVID-19 donors and healthy donors are primarily clustered by cell types, with distinct IFN signaling profiles forming subclusters within specific types (**Fig. 2D**). scMulan achieved greater consistency between IFN signaling expression and DA, with a Spearman correlation coefficient (*R*) of 0.75, surpassing the previously reported scores of 0.63 by ACR and 0.51 by CR (**Fig. 2E**). This suggests that scMulan can identify the COVID-19-associated cell state more accurately. In the analysis of individual lymphoid cell types (B cells, NK cells, CD4 T cells, and CD8 T cells), scMulan also exhibited superior performance, as evidenced by a significant increase in the area under the precision-recall curve (AUPRC) score of discriminating cells with high IFN signature by DA (**Fig. 2F and S3**, Methods). These results suggested that scMulan can well grasp the heterogeneity at finer granularity within cell types and can serve as a powerful tool for single-cell transcriptome data representation, requiring less reliance on extensive cell atlas databases and additional training processes.

### Recognizing meta-attributes of cells from gene expression with scMulan

scMulan can serve as a foundational model capable of multiple downstream tasks through generation prompted by different analytical task tokens. For instance, scMulan can characterize cells through generating meta-attributes. Here, we demonstrated this capability through cell-type annotation, the basic task in scRNA-seq data analysis. It can reflect a model’s understanding of cell identities and their gene expression signatures.

We first assessed scMulan’s ability to annotate cell types in a zero-shot manner on three datasets that had not been included in the hECA-10M data: BoneMarrow_He_2020^35^, Heart_Simonson_2023^36^, and Liver_Suo_2022^37^, which range from straightforward to complex scenarios (Methods). BoneMarrow_He_2020 comprises approximately 3,000 bone marrow cells from a single adult donor, presenting the least complexity among the three.

Heart_Simonson_2023 contains 60,345 cells from 8 human left ventricle samples, introducing challenges due to inter-individual variances. Liver_Suo_2022 contains around 140,000 liver cells from 14 fetal donors at various developmental stages, offering the most complexity among the three datasets. All three datasets underwent meticulous label alignment where all annotations from the original studies were manually mapped onto the cell types in the unified hierarchical annotation framework (uHAF) of hECA. We used accuracy, weighted precision, and weighted F1 score as evaluation metrics (Methods).

We benchmarked scMulan against CellTypist^38^, Geneformer^15^, and scGPT^16^. In this experiment, the CellTypist model was trained on a dataset subsampled from hECA-10M. Geneformer and scGPT were fine-tuned on the subsampled dataset with cell-type labels (Methods). scMulan demonstrated superior performance in all scenarios, achieving a weighted F1 score of approximately 0.9 and consistently outperforming the other models, especially in the complex datasets (**Fig. 3A**). A detailed comparison of scMulan’s annotations with the original labels confirmed its high accuracy in identifying cell types (**Fig. S4**). These results highlighted scMulan’s robust generalization ability to annotate cell types in intricate scenarios without the need for additional fine-tuning.

**Fig. 3.**
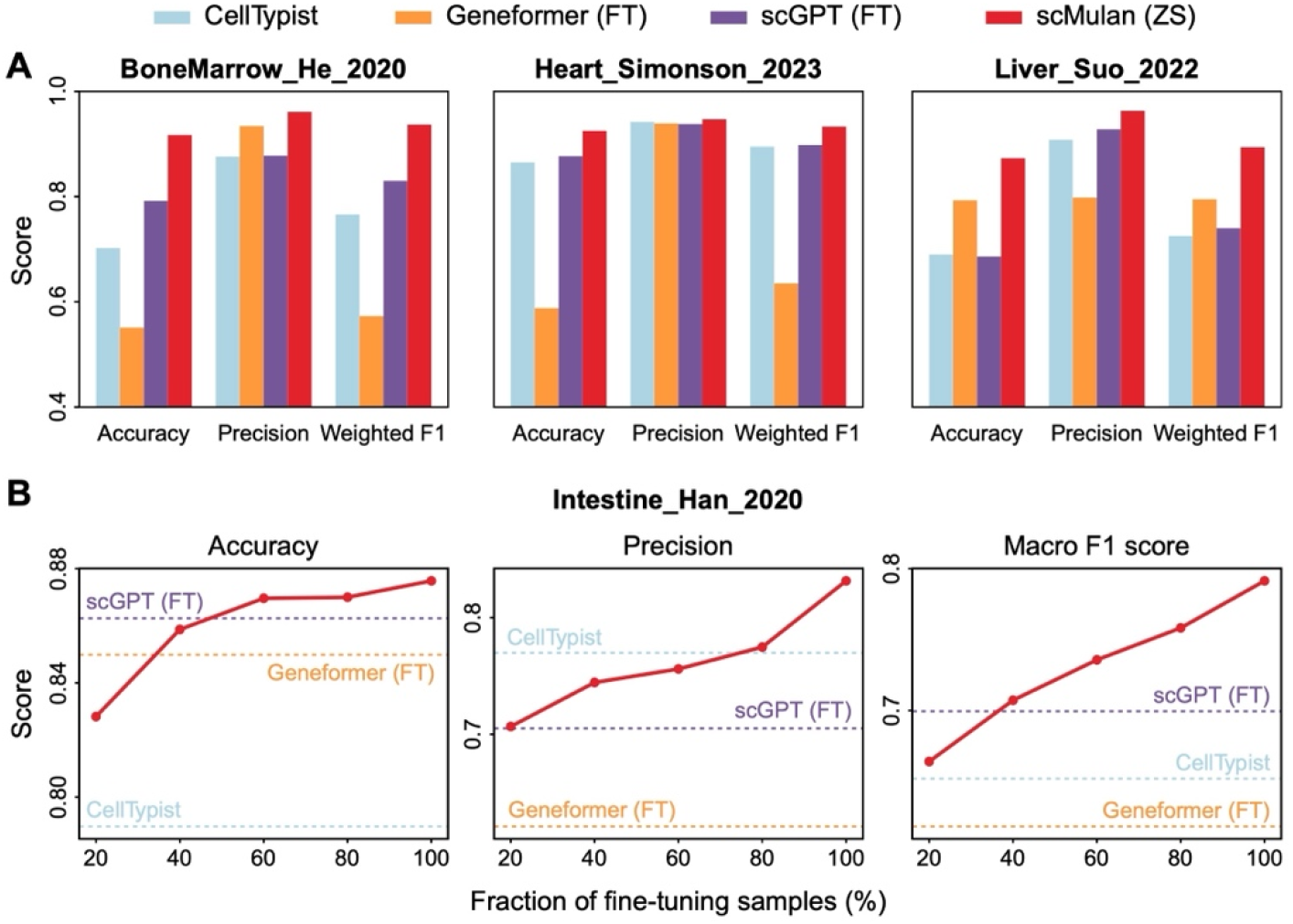
Annotating cells with scMulan. (**A**) Performance of cell type annotation for the BoneMarrow_He_2020, Heart_Simonson_2023, and Liver_Suo_2022 dataset. (**B**) Performance of cell type annotation on the Intestine_Han_2020 dataset with fine-tuning. Dashed lines represent models fine-tuned with the full training set. Red dots represent scMulan fine-tuned with different fractions of the training set. ZS, zero-shot; FT, fine-tuned.

We observed that mis-annotations of scMulan primarily occurred on cell types that were absent from the training corpus, such as secretory cells in bone marrow and endocardial cells in heart (**Fig. S4**). We thus evaluated whether fine-tuning scMulan with new data can enhance the performance, especially for organs and cell types not included in the pre-training data. We conducted an experiment focusing on the cell type annotation for intestinal, a new tissue not included in hECA-10M. We used cells of intestine in the Human Cell Landscape dataset^39^ (Intestine_Han_2020), which includes 55,214 cells across 24 cell types. We divided these cells into training and testing sets in a 9:1 ratio and incrementally fine-tuned scMulan with various fractions of cells from the training set, ranging from 20% to 100%. As a baseline, we also trained a CellTypist model and fine-tuned a scGPT and a Geneformer model on the full training set. We found that the scMulan model fine-tuned on only 40% samples of the whole training set achieved competitive performance on the testing set to the other models fine-tuned on the whole samples, and surpassed them upon fine-tuning with 60% or more of the whole samples (**Fig. 3B**). The results also indicated a consistent enhancement in performance corresponding to the increased fraction of samples used for fine-tuning the scMulan pre-trained model (**Fig. 3B**). Notably, the macro F1 score improved dramatically from 0.66 to nearly 0.80 when the fraction of cells for fine-tuning increasing from 20% to 100%, while scMulan without fine-tuning only achieved 0.25, underscoring significant performance improvements from fine-tuning on new tissues (**Fig. S5**). These results demonstrated that scMulan has superior transferability and efficiency in fine-tuning for cell characterization tasks.

To gain insight into how scMulan focuses on genes when recognizing cell types, we analyzed the genes most associated with cell type annotations, referred to as “salient genes” (Methods). Through this analysis, scMulan recovered well-established marker genes, such as *ACTA2* and *MYH11* in smooth muscle cells, *DCN* in fibroblasts, and *CD79A* in B cells^40^ (**Fig. S6**), indicating that the model has grasped key relations of marker genes and cell types. Beyond these known markers, scMulan also discovered some additional genes, suggesting novel gene expression patterns that warrant further investigation.

We also tested scMulan’s cross-species transferability for zero-shot cell type annotation using the mouse dataset, Tabula Muris^41^. We mapped mouse gene symbols to human genes using the BioMart package^42^, and then calculated the embeddings of these cells with scMulan. Results showed that the scMulan embeddings also retained the organ and cell type information of the mouse dataset with minimal batch effects (**Fig. S7A-B**). We further directly applied scMulan to the mapped gene expression data to predict cell types. Data from three tissues—liver, lung, and marrow—were tested. scMulan achieved multilabel classification accuracy of 0.97 on the liver dataset, 0.73 on the lung dataset, and 0.51 on the marrow dataset (**Fig. S7C**). These results illustrated that the information scMulan learned has strong cross-species transferability although no particular treatment has been involved for cross-species tasks in its design.

### Generating pseudo gene expression profiles of designated cells along biological processes with scMulan

Being able to create meaningful new contents is a key indicator of having mastered a language. As a generative model aiming at mastering the cell language, scMulan showed the capability of generating pseudo expression profiles of designated cells under conditions defined by meta-attributes such as age, gender, organ, and cell types (termed as meta-attribute prompts). And the generated transcriptomic profiles can be correctly identified as the cells they were designed to be by current single-cell analysis tools, passing the basic test that the created content is perceived as its intended meaning.

We first used scMulan to generate cells of different types from different organs. We combined the organs and cell types present in the hECA-10M dataset to create 140 unique pairs of organ-cell type prompts for conditional generation. Using these prompts, we generated 18,000 cells (hereafter referred to as “generated data”) and compared them with an equivalent number of cells sampled from hECA-10M (hereafter “real data”), maintaining the same cell-type proportions (Methods).

The UMAP visualization revealed similar distributions between the generated data and the real data (**Fig. 4A**), indicating that the generated data successfully reproduced the intrinsic structure and variations of the real data. We quantitatively assessed the similarity between the generated and real data with the distribution of gene expression and cell sparsity (the ratio of non-expressed genes per cell) (**Fig. 4B**). Results showed remarkable consistency in the average expression levels of all genes. We also observed that the generated data exhibited greater sparsity in gene expression compared to the real data. We analyzed that this difference in sparsity was due to scMulan’s tendency to terminate the c-sentence “prematurely”: it tends to “believe” that the number of expressed genes does not need to be too large to reproduce the desired cell type and organ source information, implying that the other genes may be irrelevant to the prompted task. This phenomenon can be used in future researches on the essential gene expressions that determine the identity and function of cells in certain tissue contexts.

**Fig. 4.**
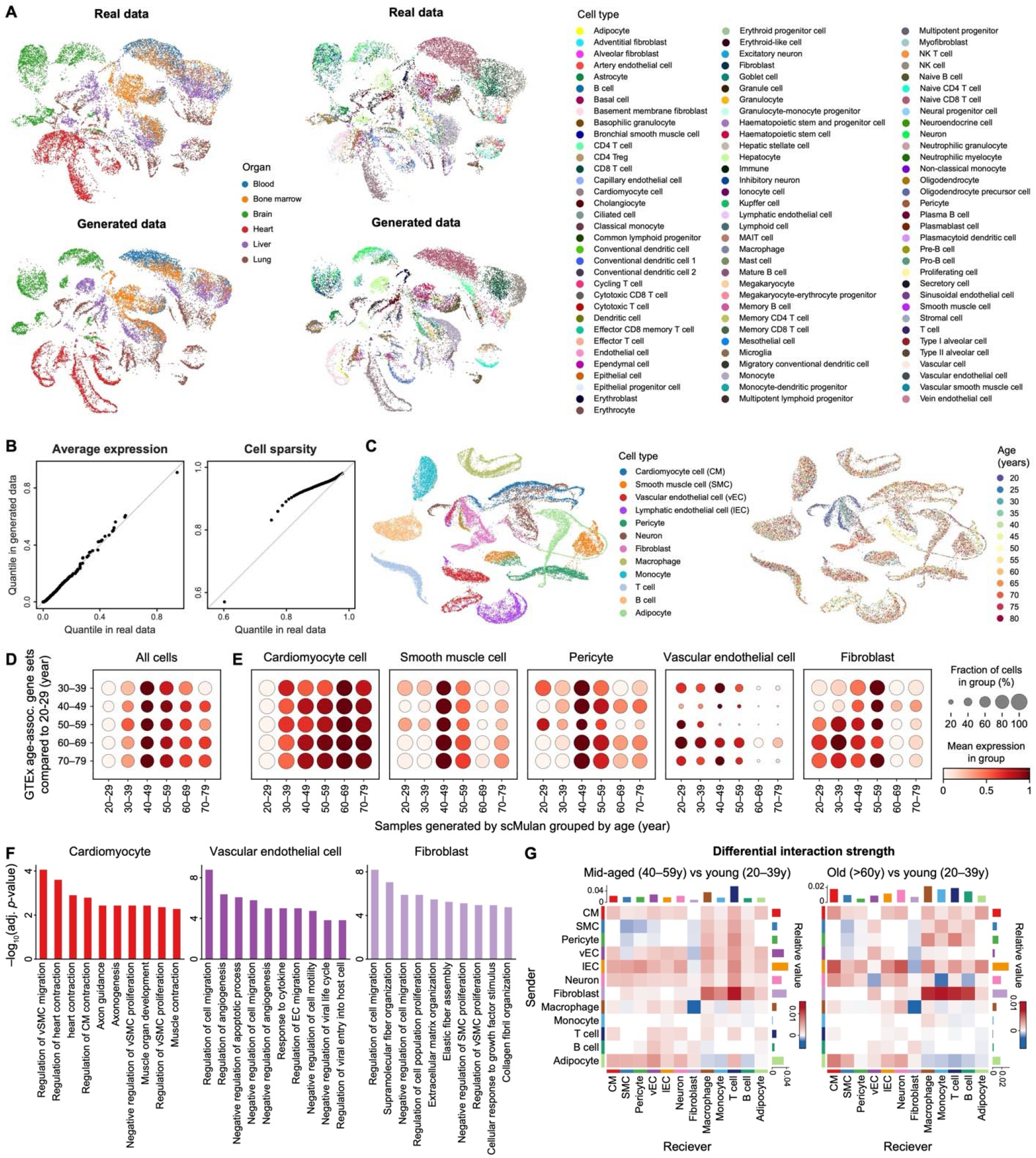
Generating pseudo gene expression profiles of designated cells with scMulan. (**A**) UMAP visualization of the real and generated hECA-10M data. Cells are colored by cell types. (**B**) The similarity between the real and generated hECA-10M data on the distribution of gene expression and cell sparsity. (**C**) UMAP visualization of the generated heart data with different donor ages. Cells are colored by cell types and donor ages. (**D**) Expression levels of the GTEx age-associated gene sets in different ages in the generated heart data. (**E**) Expression level of GTEx age signature genes in different ages in different cell types in the generated heart data. (**F**) GO enrichment analysis results of the DEGs between cells from the aged group (age over 60 years) and those from younger groups within each cell type. (**G**) Cell–cell communication networks between different age groups. CM, cardiomyocyte cell/cardiac muscle cell; SMC, smooth muscle cell; EC, endothelial cell; vSMC, vascular associated SMC; vEC, vasclular EC; lEC, lymphatic EC.

As a natural application of scMulan’s generation capability, it can be used to simulate various biological processes and investigated their differences by generating and comparing cells generated with different meta-attribute prompts. We used a case study of heart aging to showcase this capability. In this experiment, we designed meta-attribute prompts as combinations of organ, cell type, donor age, and gender. The organ was fixed as “heart”, the cell type was selected from 12 cell types common in hearts (including adipocytes, cardiomyocytes, fibroblasts, smooth muscle cells, pericytes, vascular endothelial cells, lymphatic endothelial cells, macrophages, monocytes, B cells, T cells, and neurons), the gender was chosen as either “male” or “female”, and the donor age was selected from 13 candidate values in five-year intervals from 20-to 80-years-old. We generated 100 cells for each combination of these meta-attribute tokens, resulting in a simulated dataset of 31,200 cells.

The UMAP visualization of this simulated dataset revealed clearly distinguishable patterns across cell types with slight variations in distribution along the donor ages (**Fig. 4C**). The expression profiles of marker genes (**Fig. S8A**) and the automatic annotation results by CellTypist (**Fig. S8B**) validated the reliability of the cell type variance in the generated cells.

We used the aged-associated gene sets from the GTEx project^43^ to assess the donor age in the meta-attribute prompts guided the generation. GTEx grouped heart RNA-seq expression profiles in each decade and identified highly expressed genes in each older group compared to the 20–29 age group. We regrouped the generated data accordingly and calculated the average expression of these gene sets as the aging signal. We found a consistent aging signal between the RNA-seq results in GTEx and the generated single-cell data (**Fig. 4D**). Furthermore, the generated single-cell data enabled us to examine the aging process in each cell type individually. We observed an asynchronous aging process across different cell types (**Fig. 4E**), where vascular endothelial cells showed aging signals first, followed by fibroblasts, smooth muscle cells, and pericytes, while cardiomyocytes exhibited aging signals later, particularly after age 60. This agrees well with physicians’ understanding on the aging of cardiovascular systems^44–46^.

To further explore the molecular changes during the heart aging process, we analyzed the variable genes along the aging trajectory and examined the functional differences between age groups within different cell types (Methods). We calculated the differentially expressed genes (DEGs) between cells from the aged group (age over 60 years) and those from younger groups within each cell type, and performed Gene Ontology (GO) enrichment analysis to investigate their functions (**Fig. 4F**). In aged cardiomyocytes, DEGs were primarily enriched in GO terms related to cardiac contraction, while DEGs in aged vascular endothelial cells were enriched in GO terms related to cell migration and angiogenesis. In aged fibroblasts, DEGs were enriched in GO terms related to cell migration and extracellular matrix (ECM) organization. These findings, based solely on the generated pseudo cells, are consistent with current knowledge that cardiomyocyte contractile function and endothelial cell proliferative capacity decline with age, and that changes in ECM composition and fibrosis may occur in aging hearts^47^. We also conducted a cell-cell communication analysis to investigate changes in intercellular relationships during aging. We calculated communication networks using CellChat^48^ based on ligand-receptor pairs and compared the inferred communication patterns between age groups. We observed increased communication between fibroblasts and several immune cell types in the aged heart (**Fig. 4G**), suggesting an association with inflammation, fibrosis, and fibroblast activation during heart aging. These results align with known aging-related processes in cardiac tissues^49–51^.

All these results demonstrated that the pseudo cells generated by scMulan have captured the essential mechanisms in the biological process, showing evidences of its mastering of the cell language. It opens an *in-silico* approach for studying biological processes with pseudo gene expression profiles of designated cells that may not be available or easily accessible from *in vivo* samples.

### Filling in missed cells along functional gradients with scMulan

In addition to meta-attribute prompts, scMulan also allows for pseudo cell generation using conditions of combinations of given genes and their expression levels, referred to as gene prompts. With these gene prompts, scMulan can generate the remaining genes and their expression levels in an autoregressive manner, thus creating cells of states that have not been included in the training data or are hard to be captured in real samples due to technical or ethics reasons. We demonstrated this capability by investigating how scMulan reconstructs functional gradients of cells by generating cells with gene prompts.

Vascular endothelial cells in the human heart undergo a continuous change in their expression profiles along the vasculature, from arteries to capillaries and veins, forming a functional axis^52–54^. To capture the gradients along this axis with scMulan, we first used a meta-attribute prompt where the organ was “heart” and the cell type was “vascular endothelial cell” which is a coarse cell type that didn’t lead to clear subtypes related to the location along the vasculature. We then introduced the gene prompts using two genes *GJA5* (a marker gene of the arterial endothelial cell) and *PLVAP* (as a marker gene of the venous endothelial cell)^52,53^. These two genes showed a negative correlation in real hECA-10M data (**Fig. S9A**). We set the expression levels of these two genes from 0 to 1, in increments of 0.1, respectively, as the gene prompts and generated a total of 16,600 cells (Methods).

The generated cells were clustered into three groups, and analysis using CellTypist annotations revealed that all clusters primarily consisted of endothelial cell subtypes, along with small numbers of other cell types, such as smooth muscle cells and pericytes (**Fig. S9B**). The expression levels and patterns of genes in the prompts played major roles in forming the clusters. Cells in Cluster 1 showed a continuous pattern in the expression levels of the two genes in the prompts (**Fig. 5A-B**). Cluster 2 primarily consisted of cells with relatively low but non-zero expression of *GJA5* or *PLVAP*, while Cluster 3 was dominated by cells with high expression of *PLVAP* (**Fig. S9C**). These gene expression combinations were absent from the hECA-10M pre-training data (**Fig. S9A**), suggesting that these cells might represent abnormal or rare cell states not typically observed in a healthy body. This could explain why cells in Clusters 2 and 3 received more diverse and less confident annotations from CellTypist (**Fig. S9B**), as the model was unfamiliar with these states. Additionally, cells in Cluster 2 exhibited a marked increase in mitochondrial gene expression (**Fig. S9D**), which is unusual in healthy cells^55,56^. The results demonstrated scMulan’s ability to reflect abnormal cell states when generating cells based on unrealistic or out-of-reference gene prompts.

**Fig. 5.**
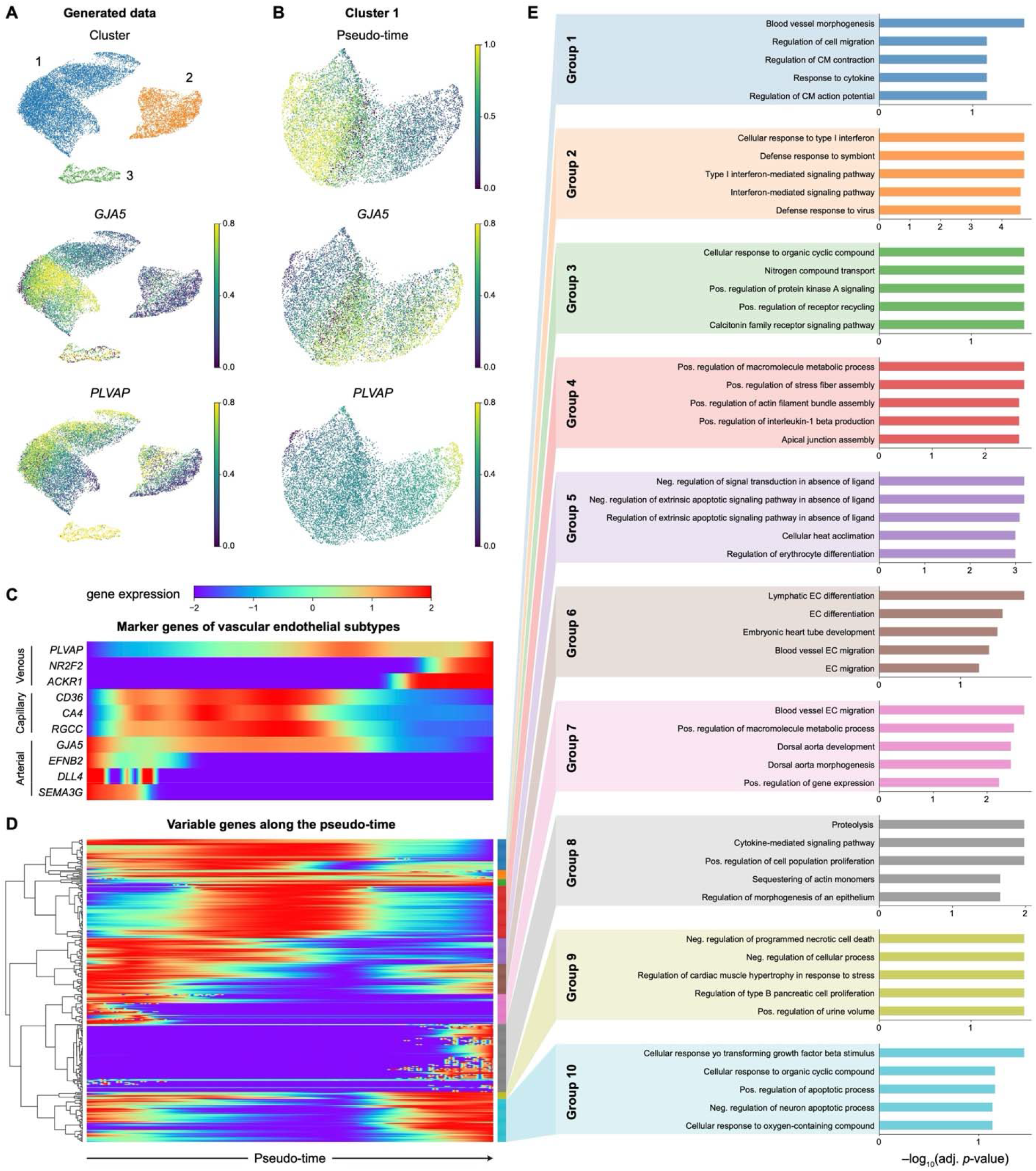
Filling in unobserved cells along functional gradients with scMulan. (**A**) UMAP visualization of the generated vascular endothelial cells. Cells are colored by clusters and gene expression levels. (**B**) UMAP visualization of Cluster 1. Cells are colored by pseudo-time and gene expression levels. (**C**) Expression of marker genes of the vascular endothelial subtypes along the pseudo-time axis of the generated cells. (**D**) Hierarchical clustering of variable genes along the pseudo-time axis of the generated cells. (**E**) GO enrichment results of different gene module groups in (D).

We focused our analysis on Cluster 1 to study the continuous transition in the generated cells. These cells were mainly predicted as capillary endothelial cells (EC1_cap, EC2_cap, and EC3_cap), arterial endothelial cells (EC5_art), and venous endothelial cells (EC6_ven) by CellTypist. We calculated pseudo-time across the transition using PAGA^57^ and confirmed a successful reproduction of the negative correlation between the two genes in the prompts (**Fig. 5B**). The marker genes for the three endothelial cell subtypes were expressed at distinct stages of the transition (**Fig. 5C**), validating that this process accurately recovered the functional gradients observed in real cells. Hierarchical clustering of variable genes along this axis identified 10 gene modules (**Fig. 5D**). GO enrichment analysis revealed that these modules were associated with biological processes such as “blood vessel morphology”, “endothelial cell migration”, and “cellular response to transforming growth factor beta stimulus”, varying along the vasculature (**Fig. 5E**). These results demonstrated that combining gene prompts with meta-attribute prompts can generate reliable data for filling in missed cells along functional gradients.

### Identifying cell fate-driver regulators with scMulan

Beyond reproducing naturally existing cell states, conditional generation by scMulan enables the prediction of gene expression profiles with gene perturbations as a built-in part of its capability, without the need for extra training or fine-tuning. The genes to be perturbed can be set in the gene prompts with desired expression levels, and the subsequent expression profiles are then generated in an autoregressive manner. This approach can simulate various gene perturbations including knockouts, knockdowns, or overexpression, and can handle any number of perturbed genes at the same time. Here, we showcased this capability with a cell-fate decision prediction task in the human hematopoiesis process.

Hematopoietic stem cells (HSCs) differentiate into several cell lineages, forming distinct branches including lymphoid cells, myeloid cells, granulocytes, megakaryocytes, and erythrocytes. Previous studies have identified key transcription factors (TFs) that govern these differentiation pathways^58^. In this experiment, we investigated whether perturbing key TFs in scMulan could drive HSCs toward specific target cell types. We selected two TFs from the human TFs in scMulan’s gene set based on current knowledge of HSC differentiation^59,60^: *GATA1*, which drives differentiation into erythrocytes and their progenitors when overexpressed, and *SPI1*, which drives differentiation into myeloid cells, including granulocytes, monocytes and their progenitors. For the metadata prompts, we set the organ as “bone marrow” and the cell type as “hematopoietic stem cell”. For the gene prompts, we varied the expression levels of *GATA1* and *SPI1* from 0 to 1, respectively, generating 100 pseudo cells for each prompt combination. Additionally, we generated 100 pseudo-HSCs without any gene prompts as the wild-type (WT) group.

To evaluate the generated perturbed cells, we collected about 9,000 hematopoietic cells from a single sample as the real cells to minimize batch effects (the HSC_Ainciburu_2023 dataset)^61^. The UMAP visualization of this dataset revealed clear differentiation branches (**Fig. 6A**). By mapping the generated cells onto this embedding space, we observed that the WT pseudo-HSCs (generated without gene prompts) aligned with real HSCs, while the perturbed pseudo cells gradually moved along the differentiation branches as the expression levels of *GATA1* and *SPI1* increased (**Fig. 6B**). At lower levels of overexpression, the perturbed cells remained near the HSC cluster, indicating stable cell states at the beginning of differentiation. As the expression levels exceeded 5, the perturbed cells began to migrate toward terminal cell types in line with the knowledge. These results demonstrated that scMulan successfully predicts HSC differentiation outcomes in response to *GATA1* and *SPI1* overexpression, validating its capability of simulating gene perturbations *in silico* without the need of fine-tuning.

**Fig. 6.**
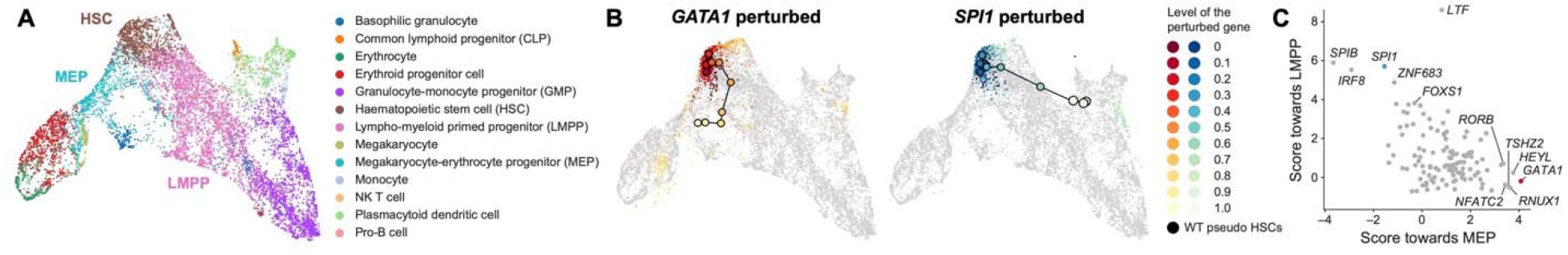
Identifying cell fate-driver regulators with scMulan. (**A**) UMAP visualization of the HSC_Ainciburu_2023 dataset. Cells are colored by cell types (**B**) Mapping the generated perturbed cells onto the embedding space of (A). Cells are colored by the level of the perturbed gene. (**C**) Relationship between the score toward MEP and the score toward LMPP for each TF.

Furthermore, scMulan can quantitatively assess the impact of various gene perturbations, offering a powerful tool for identifying key genes involved in cell fate decisions. In this study, we used UMAP to project cell embeddings and calculated the vector in the embedding space from the generated wild-type HSCs to the perturbed cells, as well as from real HSCs to their terminal cell types. We then defined a score based on the average cosine similarity between the two vectors (Methods). We calculated these scores for two critical differentiation branches: Megakaryocyte-erythrocyte progenitor (MEP) and Lympho-myeloid primed progenitor (LMPP), both of which are cell types next to HSCs on two branches in the UMAP visualization. We screened all 121 human TFs in scMulan’s gene set for single-gene overexpression and ranked them based on their scores.

We found that *GATA1* ranked first in driving differentiation toward MEP, while *SPI1* ranked third in promoting differentiation toward LMPP, consistent with their known biological roles (**Fig. 6C**). Notably, the scores for MEP and LMPP for these TFs exhibited a significant negative correlation, indicating that these two differentiation branches are mutually exclusive during cell fate decisions. Further examination of the top-ranked TFs revealed that many were previously reported to play significant roles in hematopoiesis. For example, *LTF* is specifically expressed in myeloid progenitors in mouse bone marrow^62^, and *SPIB*^63,64^ and *IRF8*^65^ are critical for myeloid cell differentiation. *ZNF683* is known to regulate human NK cell development^66^, while *RUNX1* is essential for MEP differentiation^67^, as its loss reduces MEP numbers and increases granulocyte-monocyte progenitors^68^. *NFATC2* was shown to promote differentiation toward MEP in this experiment, but was previously reported to negatively regulate this process^69^. The predicted roles of *HEYL* and *THSZ2* in fate decisions of HSCs remain to be validated.

Overall, scMulan provides a robust *in-silico* perturbation platform capable of predicting the effects of gene perturbations on biological processes. This method can offer valuable insights into candidate regulators that may drive cell fate decisions, enhancing our ability to model and manipulate cellular pathways.

## Discussion

Cells are complex systems whose mechanisms remain partially understood. While identifying individual rules in the “cell language” can bring insights into relevant biological processes, only the accumulation of such rules is insufficient for a systematic comprehension of cellular system. Current approaches focus on uncovering mechanisms piece by piece and the present the discoveries in descriptive texts and illustrations. There can be a long way to go before reaching a holistic understanding of the cell language. Individual rules or mechanisms can be validated with carefully designed experiments that controls all other factors. This strategy is no longer feasible when trying to validate the understanding of a holistic system. The solution lies in generation. As Richard Feynman said, “What I cannot create, I do not understand.” Being able to generate data that otherwise can only be obtained with experiments, and being able to pass “Turing-like tests” that can recognize the generated data as having their intended properties, can be a necessary although not sufficient proof of the comprehension of the complex system behind the data. To date, existing computational methods for single-cell transcriptomic analysis cannot yet generate novel single-cell transcriptomic profiles with designated attributes, indicating limited understanding of the comprehensive cellular systems.

This work is an attempt to explore this paradigm. We developed scMulan, a multi-task large pre-trained generative model to master the cell language from single-cell transcriptomic data by learning generate data. We designed c-sentences as an innovative way to unify both gene expression and other cellular attributes. This design enabled scMulan to handle both structured data (e.g., gene expression levels) and unstructured cellular metadata (e.g., cell type, donor demographics). We trained a 368M-parameter scMulan with the equivalence of 100M healthy human cells manually curated, standardized and annotated after data augmentation. Experiments show that the pre-trained scMulan can not only perform well on standard single-cell tasks like cell type prediction in a “zero-shot” manner, but also generate transcriptomic profiles of previously unseen cell types or states along differentiation trajectories or other gradients. It enables the creation of live cells from a few of their snapshots. It can also infer the critical genes whose expression may modulate cell state and fate. The promising results suggested that the model has acquired a substantial “understanding” of cellular language through the generative pre-training. When scaled up to more data in terms of the amount of samples, the coverage of developmental, physiological and pathological conditions, and the richness of attributes, the framework can eventually enable the construction of virtual body and biological processes with combination of real and generated “digital cells”, which can provide a powerful virtual digital life system for all types of virtual experiments.

The success of LLMs in natural language processing has demonstrated the power of generative AI to “master” complex semantics, grammar, and logical relationships of the human language. The experiments in this study, albeit preliminary, suggests that a similar generative approach may hold promise for deciphering the cellular language. Although scMulan is still in its early stages, its ability to generalize across diverse cell types and experimental conditions offers an exciting glimpse of what might be achievable through such generative LCMs with more data and more powerful models are available. These capabilities can be extended further by incorporating additional meta-attributes such as genetic variants or disease states, broadening the scope for building even more comprehensive models. Transcriptomics represents only part of the cell language. Extending scMulan to incorporate other omics data could deepen its understanding of cellular systems, which may require additional tokenization strategies and new model architectures to enable effective cross-modal learning.

The current version of scMulan has some limitations that should be addressed in the future. One technical limitation is the restriction to 2,000 highly variable genes due to computational considerations, which could limit its applicability on some biological problems. For applications that require a broader understanding of gene interactions, further fine-tuning on datasets with a wider range of genes will be necessary. As a remedy for this shortage, we designed scMulan wo allow users to customize their own gene sets of interest and train with their own data. Future work will include training a version of scMulan with the entire gene set. Another technical constrain is that the current tokenization method divides gene expression levels into 10 bins, which may miss some subtle but significant changes in gene expression. Increasing the number of bins or using the higher-resolution strategies of tokenization like in scFoundation^17^ could enhance the model’s accuracy. But such improvements all cause dramatic increase in the number of learnable parameters as well as the amount of data needed for pre-training, which imply higher computational cost. Building models that can balance the cost and effectiveness is still a challenge for all LCMs.

Another challenge is the understanding or interpretation of what scMulan has learned from the massive scRNA-seq data, although it has behaved like having mastered quite a lot of the complex cell language. Similar challenges also exist for other LCMs, other large AI models for computational biology, and the more mature field of LLMs^70^. It is arguable that both LLMs and LCMs remain largely “black boxes”. A related argument is that benchmarking tasks that can be well validated with biological experiments are crucial for AI models^71^. This is very true but may not always be feasible, given the complexity of biological systems and limitations in accessible techniques to observe the full situation of cells in many biological processes. As we strive toward concise and explicit explanations and validations of cellular mechanisms, it’s worth recognizing the possibility that the complexity of cells may exceed the explanatory power of existing reductionist frameworks and the qualitative way of presenting biological mechanisms. Adopting large-scale AI models that learn to generate rather than to dissect may represent a viable approach for mastering the complexity of cellular languages. AI models on natural languages and images had endured a long period of being naïve in many aspects before their recent great success. The promising capability exhibited by scMulan suggested that the community should embrace such attempts toward possible transformative approaches even if they may still be in infancy.

## Methods

### Defining c-sentences

We conceptualized the language of each cell as a “cell sentence”, comprising both gene expression and other meta-attributes, which function as “words” (**Fig. 1A**). In a c-sentence, a word is a 2-tuple comprising of an attribute and a corresponding value. There are four kinds of words in a c-sentence: word-of-gene (*W*_*G*_), word-of-meta-attributes (*W*_*M*_), word-of-task (*W*_*T*_), and word-of-special-token (*W*_*S*_). Each *W*_*G*_ defines a pair of a gene and its corresponding expression level. For instance, a gene *CD3D*, with an expression level of 0.2, would be represented as (*CD3D*, 0.2). For all other kinds of words, we assigned a value of zero as a placeholder. *W*_*S*_ represent special terms of a c-sentence, such as “#E#” which represents the end of a c-sentence. Here we provide an example of a c-sentence of a cardiomyocyte from the heart: “(Heart, 0), (Cardiomyocyte cell, 0), (A2M, 0.2), (ZNF385B, 0.6), …, (CD83, 0.1), (#E#,0)”.

### The architecture of scMulan

We build scMulan upon a Transformer decoder-only architecture^22^ inspired by OpenAI’s GPT models, with key adaptations tailored to the unique characteristics of single-cell data. These modifications encompass three primary components: dual embedding layers, attribute and value prediction heads, and flexible generation (**Fig. S1A**).

The dual embedding layers project input attributes and values into high-dimensional embeddings of the same dimension using an attribute embedding layer and a value embedding layer, respectively (**Fig. S1A**). The resulting attribute and value embeddings are then summed and serve as inputs to the Transformer layers.

To handle output generation, particularly for gene expression values, scMulan incorporates a unique value prediction head in addition to the original attribute prediction head (**Fig. S1A**). The value prediction head decodes logits from the final Transformer layer to predict gene expression levels. These dual embedding layers and the value prediction head allow scMulan to process gene expression levels at both the input and output stages effectively.

Furthermore, recognizing that genes in single-cell data do not inherently follow a fixed sequence, we eliminated the traditional position-aware design used in LLMs. For input words, positional encoding is omitted to preserve the position-insensitive mechanism of self-attention. During output generation, scMulan predicts all possible attributes and values of a c-sentence, conditioned on the provided input words at each time step during training (**Fig. S1A**). During inference, it generates one attribute at a time, sampled from the distribution of logits from the attribute prediction head. Although the input sequence respects the positional order of a c-sentence, we introduced random shuffling of genes in c-sentences to enhance robustness and reduce reliance on input order. This approach, termed flexible generation, introduces variability in the input and output handling, ensuring scMulan performs robustly across diverse data configurations.

### hECA-10M dataset for pre-training and validation

We curated a subset from hECA^24^ for training scMulan, named hECA-10M. The hECA-10M dataset contains over 10 million high-quality single-cell transcriptomic profiles derived from key human organs and tissues, including the heart, brain, lung, liver, bone marrow, blood, and thymus (**Fig. S1B**). Each cell in hECA-10M is characterized by the expression of 42,117 genes, accompanied by metadata capturing attributes such as the organ of origin, specific region within the organ, donor age, donor gender, sequencing technology, and cell type. This rich and diverse dataset enables the design and compilation of metadata and gene expression profiles for multiple pre-training tasks, without requiring any supervision or additional labeling.

### Pre-training strategies of scMulan

The pre-training of scMulan leverages a cell language modeling schema, which is distinct from the sequential dependency seen in natural language processing by LLMs. In traditional natural language models, the joint probability distribution of a sentence *x*, consisting of variable-length sequences of symbols (*s*_1_, *s*_2_,…, *s*_*n*_), is calculated by multiplying the conditional probabilities:

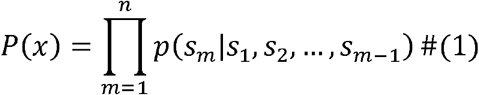

These models aim to maximize the log-likelihood of this probability distribution by estimating *p*(*s*_*m*_|*s*_1_, *s*_2_,…, *s*_*m*-1_), which is the probability of a symbol based on its preceding symbols to predict the next symbol in the sequence.

In contrast, scMulan addresses the non-sequential nature of gene orders in c-sentences. Instead of predicting the next gene, for a c-sentence *c* = {(*a*_1_, *ν*_1_), (*a*_2_, *ν*_2_),…, (*a*_*n*_, *ν*_*n*_)}, where *a*_*i*_ is the *i*-th attribute (either a gene or a meta-attribute) and *v*_*i*_ is the corresponding value, scMulan aims to estimate *p*({*a*_rest_}| {(*a*_obs_, *ν*_obs_)}, where {(*a*_obs_, *ν*_obs_)}{(*a*_1_, *ν*_1_), (*a*_2_, *ν*_2_),…, (*a*_*i*_, *ν*_*i*_)}is the set of observed attributes and their corresponding values, and is the set of rest attributes not yet conditioned upon. The goal is thus to minimize the negative log-likelihood

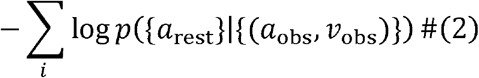

This approach can make efficient use of the self-attention computation of the Transformer decoder. This differs significantly from the masked language modeling methods used in other models like Geneformer^15^, scGPT^16^, and scFoundation^17^, which focus on modeling expressions for given genes.

Moreover, to enhance robustness and address the order insensitivity of input genes, gene entities within c-sentences are randomly shuffled during the training phase. This method ensures that the model remains effective regardless of the sequence of input data, which is critical in handling the non-sequential nature of single-cell transcriptomic data.

### Pre-training tasks of scMulan

Based on the cell language modeling schema, we can formulate different tasks as a unified generation task with the help of analytic task tokens (words-of-task) in c-sentences. We designed a set of pre-training tasks to enable multifaceted training that grasps the relationships between gene expressions and various meta-attributes. We established three primary tasks for the pre-training phase:

Conditional cell generation: As a foundational task, this involves generating genes and their corresponding expression values based on specified conditions such tissue type, cell type, donor age, sequence technology, given genes and their expression values, or any joint combination thereof. This task necessitates that scMulan grasps the intricate relationships interlinking genes and the meta-attribute conditions. In the training process, scMulan is set to condition on metadata terms only.

Hierarchical cell type annotation: This task involves generating annotations for cell types at dual hierarchical levels—both fine-grained and coarse-grained. The objective is to help the model comprehend the hierarchical structure of cell types and their relationships to gene expressions, aiding in better understanding of cellular homogeneity and heterogeneity.

Meta-attribute prediction: Demonstrated with organ/tissue prediction, this classification task requires scMulan to determine both the originating organ/tissue and its specific region from a given cell input. This task is crucial for integrating spatial variance in gene expression into the model’s understanding. Other meta-attributes also encompass important sources of cell diversity and can be employed for pre-training in the same way.

These tasks utilize a rich corpus of single-cell data formatted as a c-sentence corpus that integrates genes, meta-attributes, and tasks. The overarching goal is to optimize both subsequent attribute prediction and value prediction. The optimization of subsequent attribute prediction is to optimize Eq. 2, and the loss defined as:

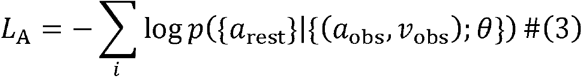

where the conditional probability *p* is modeled by scMulan’s attribute prediction head (*H*_*A*_) parameterized by *θ*.

For value prediction, we employ a mean square error (MSE) objective:

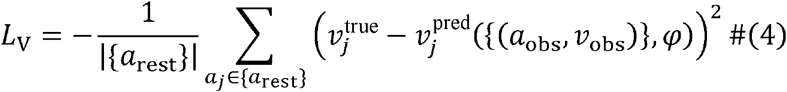

Where 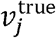 denotes the true value of attribute *a*_*j*_,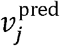 represents the predicted value from scMulan’s value prediction head (*H*_*V*_) given the observed attributes and their values {(*a*_obs_, *ν*_obs_)} with parameter *φ*.

The attribute and value prediction heads *H*_*A*_ and *H*_*V*_ leverage shared hidden embeddings derived from a *n*-layer Transformer decoder:

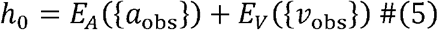

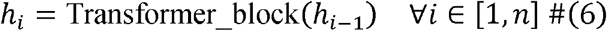

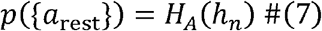

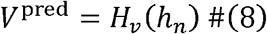

where *E*_*A*_ and *E*_*V*_ are the attribute and value embedding layers, respectively, *H*_*A*_ and *H*_*V*_ are implemented as multi-layer perceptrons (MLPs), and *V*^pred^ is the corresponding values of{*a*_*rest*_}.

We optimize the compound objective with a weighting factor λ:

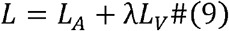

The parameters of scMulan are trained via adaptive gradient algorithms.

### Benchmarking on data integration

We evaluated how scMulan learns the contextual information of a c-sentence by benchmarking the performance of its zero-shot cell embeddings in a data integration task. The experiments were conducted on two datasets: Lung_Luecken_2022 and COVID-19_Lotfollahi_2022. The Lung_Luecken_2022 dataset, sourced from the published data integration benchmark scIB^31^, contains 32,472 cells from 3 batches across 2 sampling locations and 17 cell types. The COVID-19_Lotfollahi_2022 dataset, provided by scArches^32^, includes 274,346 cells from 18 batches, capturing diverse COVID-19 severities. For both datasets, we standardized gene symbols using the GeneSymbolUniform toolkit from hECA^24^, aligning them to 42,117 unique genes. The data was normalized to a total count of 10,000 and then log-transformed. To ensure a fair comparison across methods, we selected the 2,000 genes that were employed for training scMulan as input features for all methods, termed as the scMulan-2K gene set.

We tested task-specific integration methods, including BBKNN^25^, Harmony^26^, Seurat v3^27^, FASTMNN^28^, scVI^29^, and scANVI^30^, as well as foundation models such as GeneFormer^15^, scGPT^16^, scFoundation^17^, and our scMulan. Among the task-specific methods, FastMNN and Seurat were implemented in R using the Seurat and SeuratWrappers package, while others were implemented in Python. Default parameters were used for all methods. The scANVI models were trained based on the trained scVI models and the additional cell-type annotation inputs.

For Geneformer and scFoundation, contextual cell embeddings were extracted according to their tutorials without further fine-tuning on the integration task. For scGPT, experiments were conducted in both zero-shot and fine-tuning paradigms. In the zero-shot paradigm, the contextual embedding of the CLS token was used as the cell embedding for evaluation. In the fine-tuning paradigm, batch information was provided to the scGPT model, and the model was fine-tuned using adversarial learning for 15 epochs. For scMulan, we set a c-sentence composed of a cell’s gene expression profile followed by the task token “<cell type generation>” as inputs, and extracted the representation from the final Transformer layer of scMulan as the cell embedding.

To evaluate performance, we saved the embeddings from each method and calculated AvgBIO scores, defined as the average of *k*-means normalized mutual information (NMI), *k*-means Adjusted Rand Index (ARI), and Silhouette label metrics using the scIB-metric package^31^. For runtime comparison, we recorded the time taken by each method on the COVID-19_Lotfollahi_2022 dataset, with GPU-accelerated methods leveraging an NVIDIA A100-80G GPU.

### Identifying COVID-19-associated cell states

We analyzed altered cell states in a COVID-19 dataset, COVID-19_Dann_2023, derived from a previous benchmark study^33^. The dataset, constructed with data from the CELLxGENE portal in the original paper, includes 62,509 PBMCs from both healthy and COVID-19 donors, with harmonized cell-type annotations.

In the original study, COVID-19-associated cell states were defined as IFN-stimulated cells within each cell type, which served as the “ground truth” in our experiments. The IFN signature was computed using the tl.score_genes function from the SCANPY package^72^ on a predefined gene set, including *BST2, CMPK2, EIF2AK2, EPSTI1, HERC5, IFI35, IFI44L, IFI6, IFIT3, ISG15, LY6E, MX1, MX2, OAS1, OAS2, PARP9, PLSCR1, SAMD9, SAMD9L, SP110, STAT1, TRIM22, UBE2L6, XAF1*, and *IRF7*.

To identify COVID-19-associated cells, the study constructed neighborhoods in the cell embedding space and tested differential abundance (DA) of cells from healthy and COVID-19 groups using Milo^34^. Neighborhoods with significantly higher abundances of cells from COVID-19 samples were identified as COVID-19-associated cell states. We evaluated three types of cell representation paradigms:

- The control reference (CR) design (from the original study): Healthy and COVID-19 groups were treated as separate batches and integrated using scVI^29^ to obtain joint embeddings.
- An atlas to control reference (ACR) design (from the original study): Both groups were mapped to an atlas dataset (provided by the original study) using a pre-trained scArches model^32^ for calculating cell embeddings.
- scMulan-based design: Cell embeddings were directly derived using scMulan, following the same integration method as in the data integration benchmarking experiment.

We implemented the CR and ACR designs using the original codebase (https://github.com/MarioniLab/oor_design_reproducibility) and reproduced the reported results. For downstream analysis, we constructed a *k*-nearest neighbor (*k*=100) graph for cells in each type of representations and conducted DA tests using the Milopy package^34^. Neighborhoods enriched with COVID-19 cells (defined as a positive log-transformed fold change of COVID-19 cells over healthy cells and a spatial FDR < 10% in the DA test) were identified as COVID-19-associated states.

To evaluate the identification results, we reported two metrics from the original study, Spearman’s correlation coefficient (SCC) and precision-recall curve (PRC) analysis. SCC was calculated between the mean IFN signature and the log-transformed fold change in all neighborhoods. A higher SCC indicates better consistency with the ground truth and superior cell representation. Besides, to evaluate the ability of different representations to detect COVID-19-associated cell states, we generated PRCs using various log-transformed fold change thresholds. The area under the PRC (AUPRC) was computed for the top 10% IFN-high neighborhoods. A higher AUPRC indicates better performance of the representation in accurately identifying COVID-19-associated cell states with fewer false positives.

### Benchmarking on cell type annotation

To comprehensively evaluate the performance of foundation models pre-trained on a multi-organ corpus, we prepared four benchmark datasets from different organs for cell type annotation. The BoneMarrow_He_2020 dataset includes 3,230 cells from an adult bone marrow sample in the Adult Human Cell Atlas (AHCA)^35^. The Heart_Simonson_2023 dataset was constructed by selecting cells from control group samples in a study on end-stage heart failure^36^. We selected healthy donors (P1452, P1650, P1690, P1716, P1739, P1763, P1785, and P1801) mentioned in the original study, resulting in 60,345 nuclei. The Liver_Suo_2022 dataset was created by selecting liver cells from a multi-organ atlas studying the development of the human immune system^37^. We extracted 137,181 liver cells from 13 fetal livers. The Intestine_Han_2020 dataset was derived from the Human Cell Landscape (HCL)^39^. This dataset includes 55,214 intestine cells from fetal and adult donors, covering different regions of the intestine. The data were accessed via our published hECA^24^, where samples were merged and cell type annotations were harmonized. It should be noticed that none of these datasets are included in the hECA-10M dataset.

We also standardized gene symbols of these datasets to 42,117 genes with GeneSymbolUniform^24^. All datasets underwent preprocessing with total-count normalization to 10,000 and log-transformation. To ensure a fair comparison across methods, we selected the scMulan-2K gene set as input features for further analysis. We manually mapped the original annotations to the unified hierarchical annotation framework (uHAF)^24^. The detailed mapping relationships are available in our codebase.

We evaluated scMulan’s zero-shot annotation performance on BoneMarrow_He_2020, Heart_Simonson_2023, and Liver_Suo_2022—organs included in scMulan’s pre-training. For comparison, we fine-tuned scGPT and Geneformer on cell type annotation tasks, following their tutorials and using default hyperparameters. To map the output of fine-tuned models to uHAF cell types, we randomly selected 200,000 cells from hECA-10M, splitting them 80:20 for fine-tuning and validation. We also trained a CellTypist model on this dataset. scMulan predicted fine-grained cell types by prompting with the input cell’s gene expression, followed by a “<cell type generation>” task token.

For Intestine_Han_2020, we split the dataset into training and test sets at a 9:1 ratio. Geneformer and scGPT were fine-tuned, and a CellTypist model was trained on the training set. Since scMulan was not pre-trained on intestine data, we assessed its transfer learning performance by fine-tuning on different fractions of the training set (20%, 40%, 60%, 80%, and 100%). We used the “<cell type generation>” task token to fine-tune the scMulan model with the same approach during pre-training. All models were then used to predict cell types on the test set.

We evaluated model performance using accuracy, precision, and F1-score, calculated with the scikit-learn package^73^. For the zero-shot annotation experiment, where models output cell types across multiple organs, weighted-F1 scores were used to account for class imbalance by weighting scores by the number of cells per type. In contrast, for the intestine experiment, where predictions and ground truths were context-specific, we used macro-F1 scores. Additionally, predictions were relaxed to count as correct if they matched a subclass of the true label. For example, a cell predicted as a CD8 T cell was deemed correct if the true label was “T cell” or “Lymphoid cell”.

### Saliency gene identification

We implemented gradient-based explanation techniques to analyze how each input feature, particularly gene, influences the model’s cell type predictions. This process involved calculating the partial derivatives of the predicted cell types with respect to the embeddings of each gene, a technique based on the Gradient X Input method^74^, which allows us to compute attribution scores for each gene in a c-sentence under the cell type prediction task.

We then pinpointed the top five genes per cell based on their attribution scores, which reflect their importance or relevance in determining cell types. To further distill this information, we identified the ten most frequently occurring genes across these top-ranking attribution scores for each cell type. These genes are considered “saliency genes” because they potentially act as markers, distinguishing one cell type from another.

### Generating cells with prompts

To evaluate scMulan’s capacity for conditional generation, we established various organ and cell type pairings as joint conditions under which the model can generate cell expression profiles. We chose organs including the heart, lung, brain, blood, bone marrow, and liver, and for each, we randomly generated 3,000 cells, each assigned a cell type corresponding to the particular organ. This process yielded a total of 18,000 organ-cell type pairs and their associated cells generated by scMulan. The organ-cell type pairs were converted into c-sentences, appended with the task token “gene expression generation” to direct scMulan for conditional cell generation. For example, the c-sentence “(Heart, 0), (T cell, 0), (<gene expression generation>, 0)” would prompt scMulan to generate a T cell specific to the heart.

Besides, to simulate the aging process of the heart, we generated cells using meta-attributes including organ, cell type, age, and gender. The organ was fixed as “heart”, and gender was specified as “male” or “female”. Cell types included adipocytes, cardiomyocytes, fibroblasts, smooth muscle cells, pericytes, vascular endothelial cells, lymphatic endothelial cells, macrophages, monocytes, B cells, T cells, and neurons, representing common cell types found in the human heart. To cover the entire aging spectrum of adult hearts, ages were chosen from 13 values in five-year intervals ranging from 20 to 80 years old. For example, a sample prompt “(Heart, 0), (Fibroblast, 0), (50 year, 0), (M, 0), (<gene expression generation>, 0)” would instruct the simulation to generate fibroblasts from the heart of a 50-year-old man. For each combination of these meta-attributes, we simulated 100 cells, resulting in a constructed dataset comprising 31,200 cells.

The integration of “word-of-gene” and “word-of-meta-attribute” in c-sentences empowered scMulan to generate cells with specific gene expression profiles alongside pre-defined meta-attribute conditions. Similar to conditional generation based on meta-attributes, scMulan allows for combinations of any number of gene expression conditions by incorporating corresponding “word-of-gene” prompts into the c-sentence prompt. Then scMulan could generate the remaining genes with a “<gene expression generation>” task token. We utilized this capability to simulate functional gradients in heart endothelial cells and to explore cell fate decisions in hematopoietic stem cells (HSCs).

In simulating continuous functional gradients along the vasculature in the human heart, we focused on two marker genes, *GJA5* and *PLVAP*, alongside fixed meta-attribute conditions specifying the organ as heart and the cell type as vascular endothelial cell. We varied the expression levels of these genes from 0 to 1, incrementing by 0.1. An example prompt would be “(Heart, 0), (Vascular endothelial cell, 0), (GJA5, 0.1), (PLVAP, 0.8), (<gene expression generation>,0)”. For each combination of conditions, we generated 200 cells, resulting in a dataset of 16,600 cells. To assess the impact of gene expression conditions, we additionally generated 400 cells using meta-attribute conditions alone, with a prompt structured as “(Heart, 0), (Vascular endothelial cell, 0), (<gene expression generation>,0)”.

To simulate the response of HSCs to gene perturbations, we individually perturbed one gene at a time by including the corresponding “word-of-gene” in the prompt starting with “(Bone marrow, 0), (Hematopoietic stem cell, 0)”. For each selected gene, we varied its expression from 0 to 1 in increments of 0.1 and generated 100 cells for each expression level. Specifically, we perturbed genes like *GATA1* and *SPI1* to observe their effects on HSCs. Additionally, we generated 100 pseudo HSCs without any gene prompts using the prompt “(Bone marrow, 0), (Hematopoietic stem cell, 0), (<gene expression generation>,0)”. In the study focusing on 121 human transcription factors (TFs), we set the expression level of each TF from 0.1 to 1 and generated 20 cells for each value. scMulan produced 200 cells for each TF, resulting in a dataset containing 242,000 cells.

### Analysis of the generated heart data

We analyzed the generated heart data using a standardize pipeline in the Scanpy package^72^. Due to the binned expression format of the generated data, we skipped the normalization and log-transformation steps. Principal Component Analysis (PCA) was performed, and the first 15 principal components were selected for downstream neighbor graph construction, where the parameter n_neighbors was set to 30.

To validate that the cell type information in the meta-attribute condition worked, we used a pre-trained CellTypist model to annotate the data and compared the predictions with the cell types in the meta-attribute conditions. The model, named “Healthy_Adult_Heart”, was download from https://www.celltypist.org/models. To conform to the model’s input requirements, we performed an inverse log1p transformation on the binned expression values using the numpy.expm1 function and re-normalized the total counts for each cell to 10,000. The processed data were then input into the CellTypist model, with majority_voting set to True during prediction. Subsequently, we returned to the original binned expression data for all following analyses.

For aging signature analysis, we utilized GTEx aging-associated gene sets obtained from https://maayanlab.cloud/Enrichr/geneSetLibrary?mode=text&libraryName=GTEx_Aging_Signatures_2021. These gene sets were identified by comparison between the expression profiles of tissue samples in different age groups based on the GTEx database. We selected five heart-related gene sets (GTEx Heart 20-29 vs 30-39 Down, GTEx Heart 20-29 vs 40-49 Down, GTEx Heart 20-29 vs 50-59 Down, GTEx Heart 20-29 vs 60-69 Down, and GTEx Heart 20-29 vs 70-79 Down). For each gene set, the aging signature was calculated as the sum of expression values for genes overlapping with the gene set and the scMulan-2K gene set. The generated cells were re-grouped into 10-year age intervals for visualization, matching the age groups of the gene sets.

For differential expression analysis, we grouped cells into three broader age categories (20–39 years, 40–59 years, and >60 years) within each cell type. Differentially expressed genes (DEGs) were identified using the Wilcoxon test via the scanpy.tl.rank_genes_groups function. Genes with an adjusted *p*-value < 0.05 in the >60-year group were selected for downstream Gene Ontology (GO) enrichment analysis, which was performed using the Python package gseapy^75^ with the GO_Biological_Process_2023 database^76^. We also conducted the cell–cell communication analysis with the R package CellChat^48^. We calculated ligand-receptor-based cell–cell communication for each broader age group and compared the pairwise changes in communication strength at the cell-type level.

### Analysis of the generated vascular endothelial cell data

We used pre-trained human heart CellTypist models (the same as that in the heart aging experiment) for annotation. The same inverse log1p transformation and re-normalization were conducted. Then we utilized the generated data with gene prompts for visualization and clustering. PCA was applied to binned expression data, and a neighbor graph was constructed using the first 15 principal components. Clustering was performed using the Leiden algorithm with a resolution parameter of 0.1. In the visualization of UMAP, we calculated mitochondrial gene signatures using scanpy.pp.calculate_qc_metrics, focusing on the percentage of mitochondrial gene expression within the scMulan-2K gene set.

In downstream analysis, we focused on a cluster showing low mitochondrial gene signatures and clear trajectory in visualization. Subsequently, we repeated the PCA, neighbor graph construction, and UMAP visualization steps on cells in this cluster. Using the PAGA algorithm^57^ in Scanpy, we constructed a trajectory of the functional axis of human heart vasculature and calculated pseudo-time along this trajectory.

Cells were ordered based on pseudo-time values, and gene expression values were smoothed using Locally Estimated Scatterplot Smoothing (LOWESS) from the Python statsmodels package^77^. Genes with zero variance along the trajectory or failed LOWESS convergence were filtered out, leaving 303 genes for downstream clustering and visualization. To improve visualization, we scaled the expression values of each gene to [–2, 2].

To visualize gene expression changes along the functional axis, we used seaborn.clustermap^78^ with metric = ‘cosine’ and method = ‘complete’, extracting a dendrogram to cluster genes into 10 functional groups using the fcluster function from the scipy package^79^. Each gene cluster underwent GO enrichment analysis using the gseapy package and the GO_Biological_Process_2023 database.

### Analysis of the generated hematopoiesis data

To assess the impact of gene perturbations on HSCs, we utilized the HSC_Ainciburu_2023 dataset^61^, focusing on cells sampled from a 19-year-old donor to minimize batch effects. The dataset was standardized to 42,117 gene symbols using GeneSymbolUniform^24^, and we extracted the scMulan-2K gene set for binned expression transformation in scMulan preprocessing. Subsequently, PCA and UMAP models were trained using the scikit-learn package^73^ for dimensional reduction, and these models were saved for future use. To facilitate comparison with generated cells, original cell types were manually mapped onto uHAF^24^.

In experiments involving *GATA1* and *SPI1* gene perturbations, we mapped generated data onto the reference dataset using the pre-trained models. Centroids of cells with the same *GATA1* (or *SPI1*) expression level were calculated and plotted in ascending order to illustrate the differentiation process characterized by a continuous increase in the perturbed gene’s expression.

In the screening of fate-decision TFs, we defined the gene prompt score based on its ability to drive HSC differentiation towards specific branches:

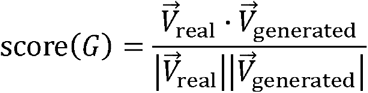

Here 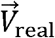 represented the vector from the centroid of HSCs to the centroid of the target cell type in the UMAP space of the real data, and 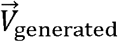represented that in the generated data with gene prompt *G*. The score was defined as the cosine similarity between the two vectors. To identify key TFs, we calculated the average score across prompts with expression levels ranging from 0.1 to 1 along the differentiation trajectories toward the Megakaryocyte-Erythrocyte Progenitor (MEP) or toward the Lympho-Myeloid Primed Progenitor (LMPP), respectively.

## Acknowledgements

The work is supported in part of National Natural Science Foundation of China (62250005, 62373210, 62433001, and 92470105), The National Key R&D Program of China (2021YFF1200900), and The Tsinghua-Fuzhou Institute for Data Technology.

## Author contributions

Y.C., H.B., L.W., and X.Z. conceived the project. Y.C., H.B., and X.D. designed the model architecture. Y.C., H.B., and J.J. designed the experiments on generated data. Y.C., H.B., Y.L., Y.Z., and X.W. generated the data using scMulan and analyzed saliency genes. Y.C. and H.B. performed data analysis. C.L., E.L., C.X., and M.H. provided valuable suggestion and assistance on model design and data analysis. Y.C., H.B., L.W., and X.Z. wrote the manuscript. X.Z. and L.W. supervised the project.

## Competing interests

The authors declare that they have no competing interests.

## Data availability

The hECA-10M dataset can be downloaded from https://doi.org/10.5281/zenodo.14209941. The Lung_Luecken_2022 dataset was downloaded from https://doi.org/10.6084/m9.figshare.12420968. The COVID-19_Lotfollahi_2022 dataset was downloaded from https://github.com/theislab/scarches-reproducibility. The COVID-19_Dann_2023 dataset was downloaded from https://doi.org/10.6084/m9.figshare.21456645. The BoneMarrow_He_2020 dataset is accessible via the Gene Expression Omnibus (GEO) database under the accession number GSE159929. The Heart_Simonson_2023 dataset was downloaded from the Broad Institute’s Single Cell Portal under the accession number SCP1849. The Liver_Suo_2022 dataset was downloaded from https://developmental.cellatlas.io/fetal-immune. The Intestine_Han_2020 dataset and the HSC_Ainciburu_2023 dataset we pre-processed can be downloaded from https://doi.org/10.5281/zenodo.14209910.

## Code Availability

The codes for experiments in this study and the scMulan model is available at https://github.com/cyxss/scMulan.

## Notes

### Competing Interest Statement

The authors have declared no competing interest.

### Summary of Updates

Results updated; author list updated; figures updated.

