## Supplementary materials for "Toward mastering the cell language by learning to generate"

### Supplemental Figures

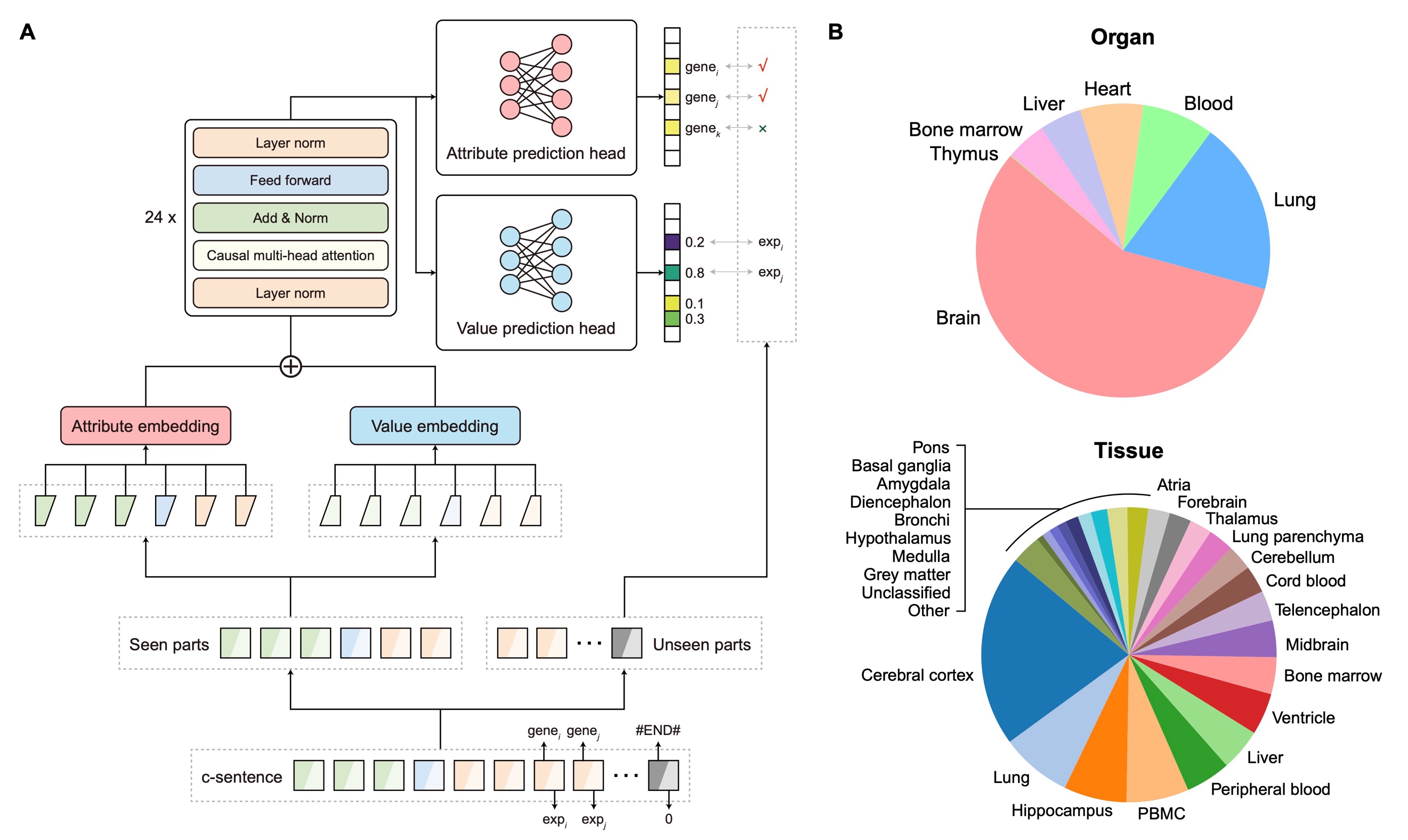

**Fig. S1. Overview of scMulan.** (**A**) The architecture of scMulan. The entity embedding and value embedding layer take the entities and their corresponding values from the c-sentences as input, respectively, and project them as the high dimensional embeddings. The transformer takes the summation of these two types of embeddings as input. scMulan has two prediction heads, one for attribute prediction, and the other for value prediction. At each time step, scMulan predicts the unseen attributes and values of the c-sentence, condition on the seen entities and values. (**B**) Overview of the hECA-10M dataset.

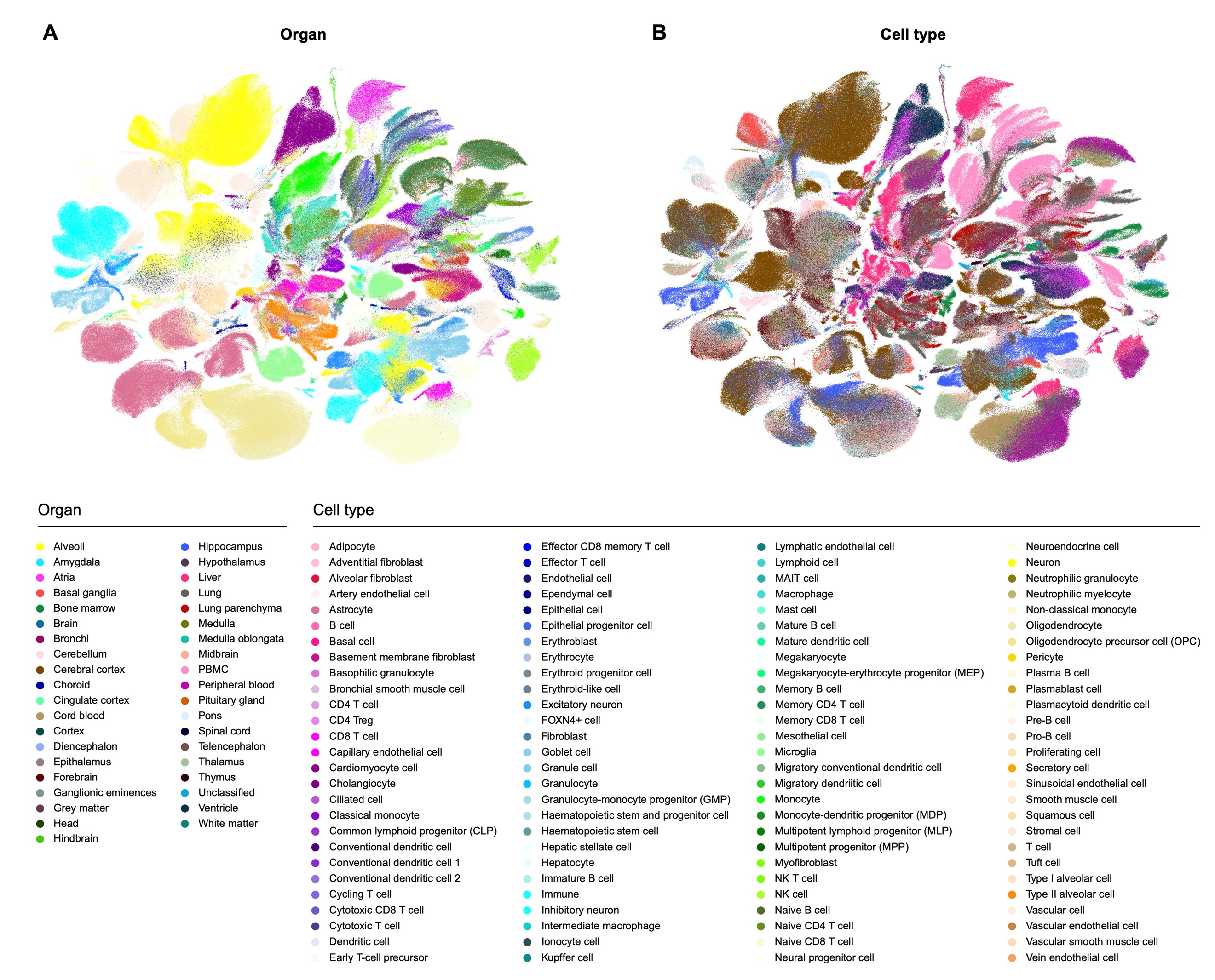

**Fig. S2. UMAP visualization of the scMulan embeddings of the validation set of hECA-10M.** Cells are colored by organs (**A**) and cell types (**B**).

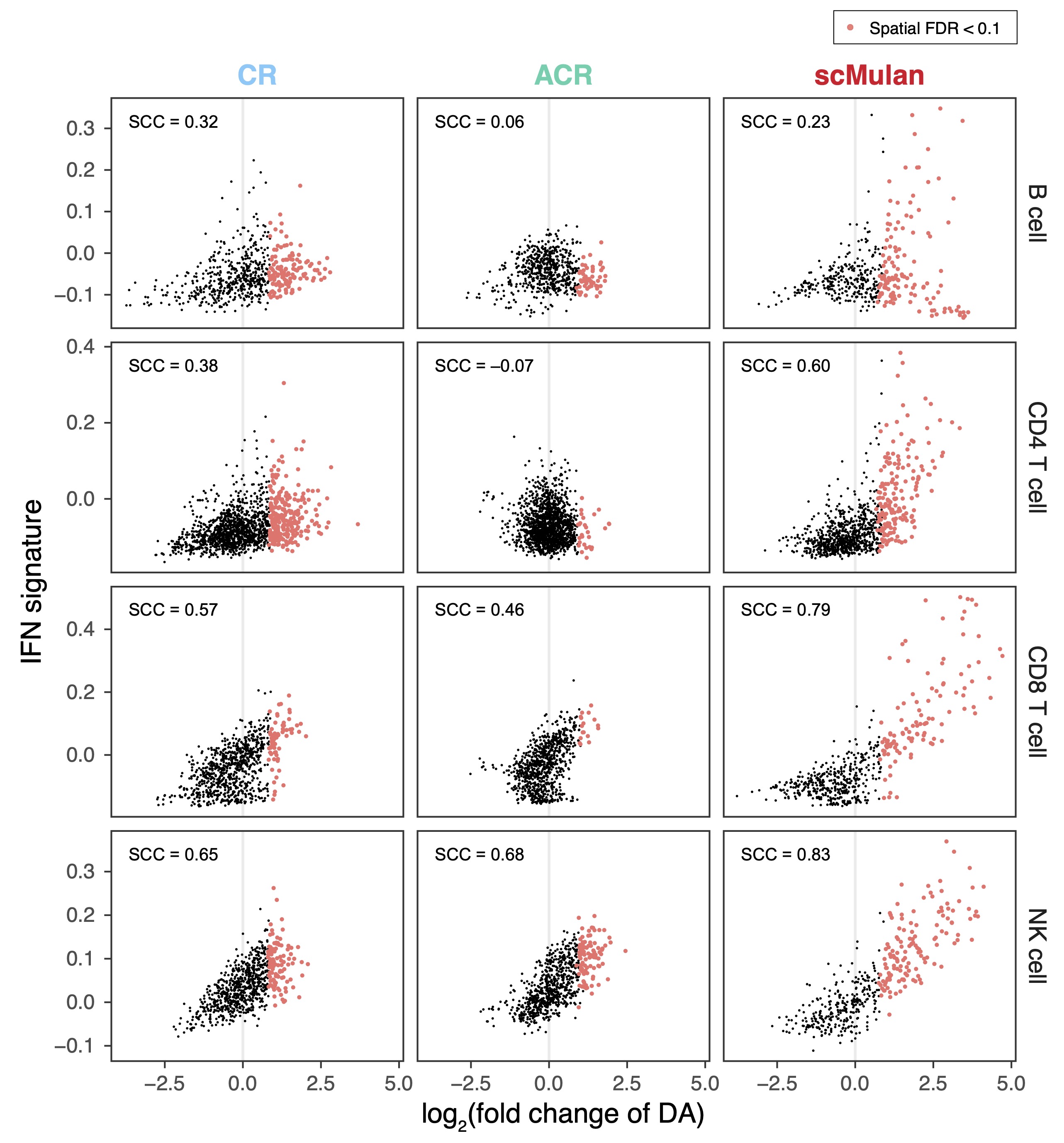

**Fig. S3. Relationships between DA and the IFN signature for different lymphoid cell types in COVID-19_Dann_2023.** Cells with neighborhoods significant enriched in COVID-19-associated cells (log_2_(fold change of DA) > 0 and spatial FDR calculated by Milo < 10%) are colored with red.

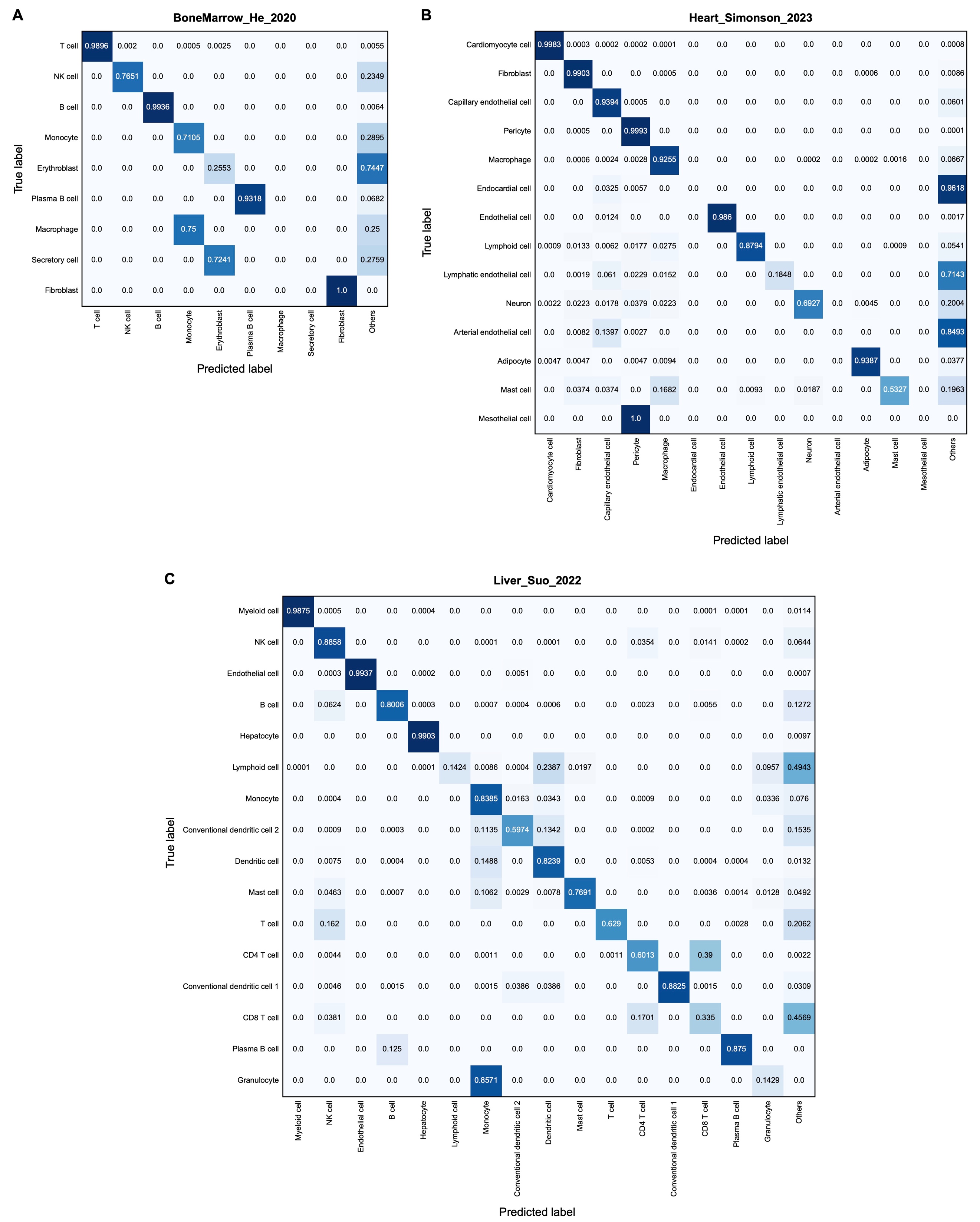

**Fig. S4. Confusion matrices between cell type annotations from the original study and those by scMulan without fine-tuning.** (**A**) The BoneMarrow_He_2020 dataset. (**B**) The Heart_Simonson_2023 dataset. (**C**) The Liver_Suo_2022 dataset.

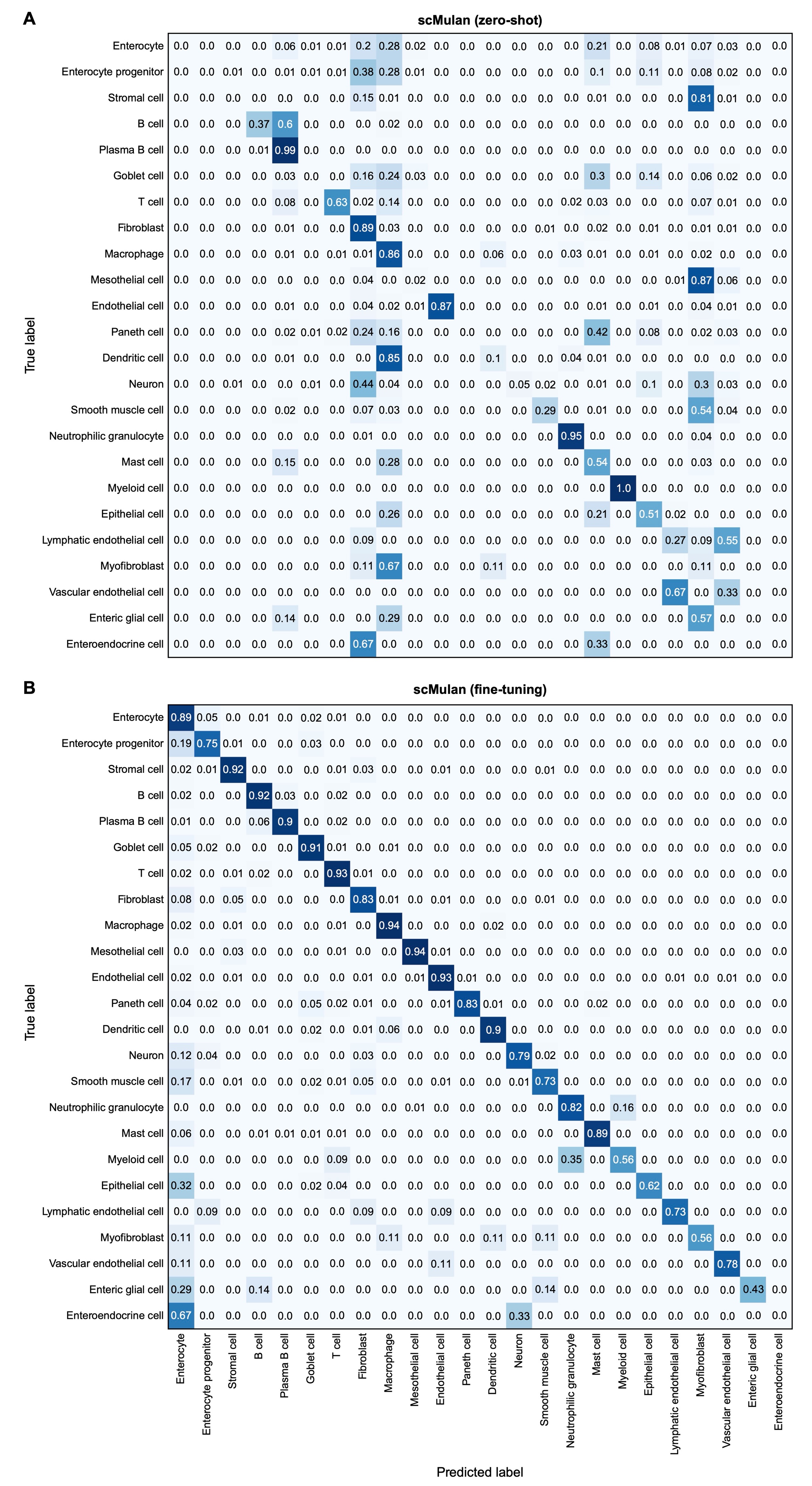

**Fig. S5. Performance of scMulan without and with fine-tuning on annotating the Intestine_Han_2020 dataset.** (**A**) Without fine-tuning. (**B**) With fine-tuning.

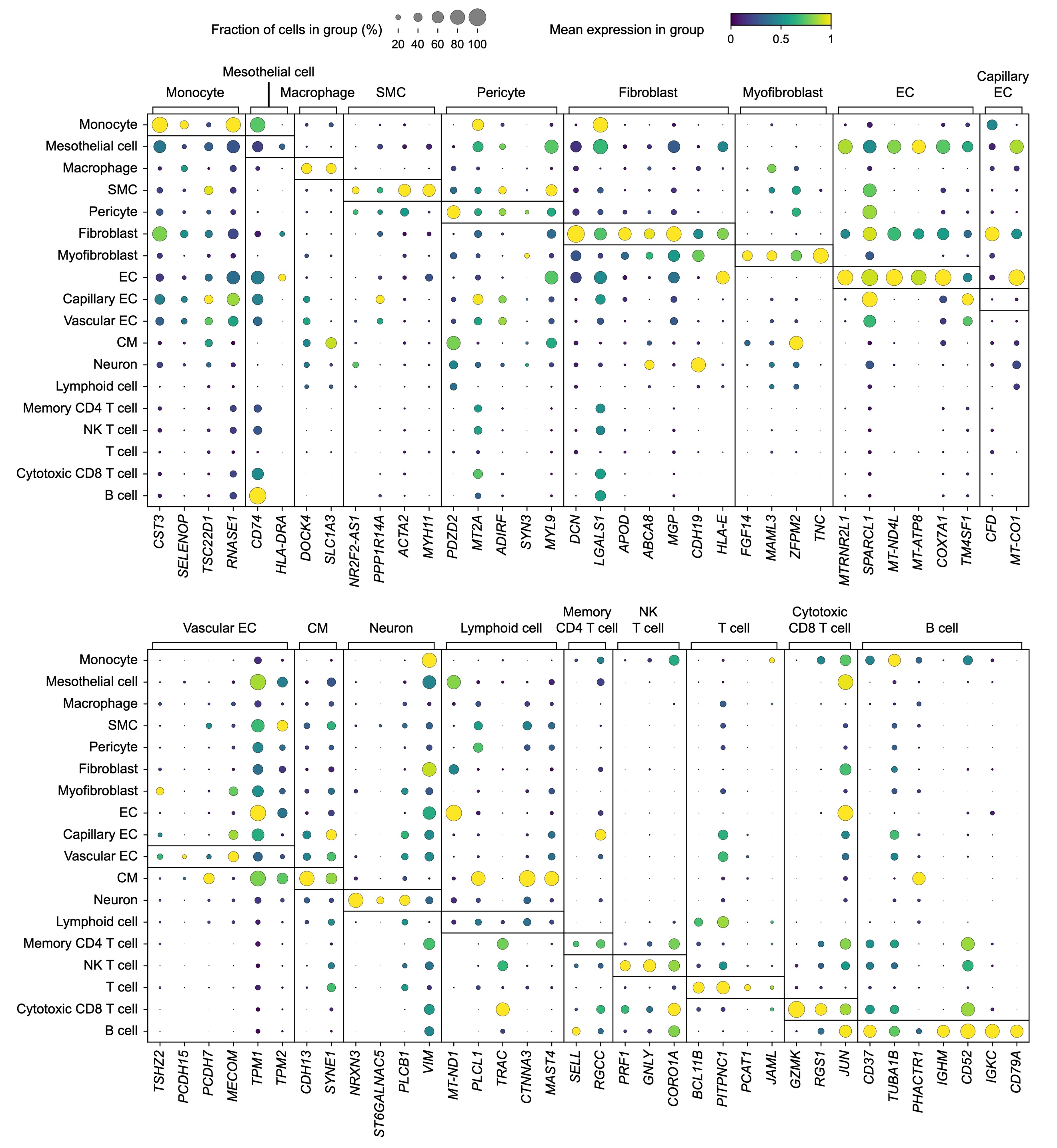

**Fig. S6. Bubble plot visualization of saliency genes extracted by scMulan in heart cells of hECA-10M.** Each row corresponds to a specific cell type. Each column represents a saliency gene, grouped by its associated cell type. we identified the ten most frequently occurring genes across the top-ranking attribution scores for each cell type as saliency genes, and genes that are salient in more than one cell type are excluded. CM, cardiomyocyte cell; SMC, smooth muscle cell; EC, endothelial cell.

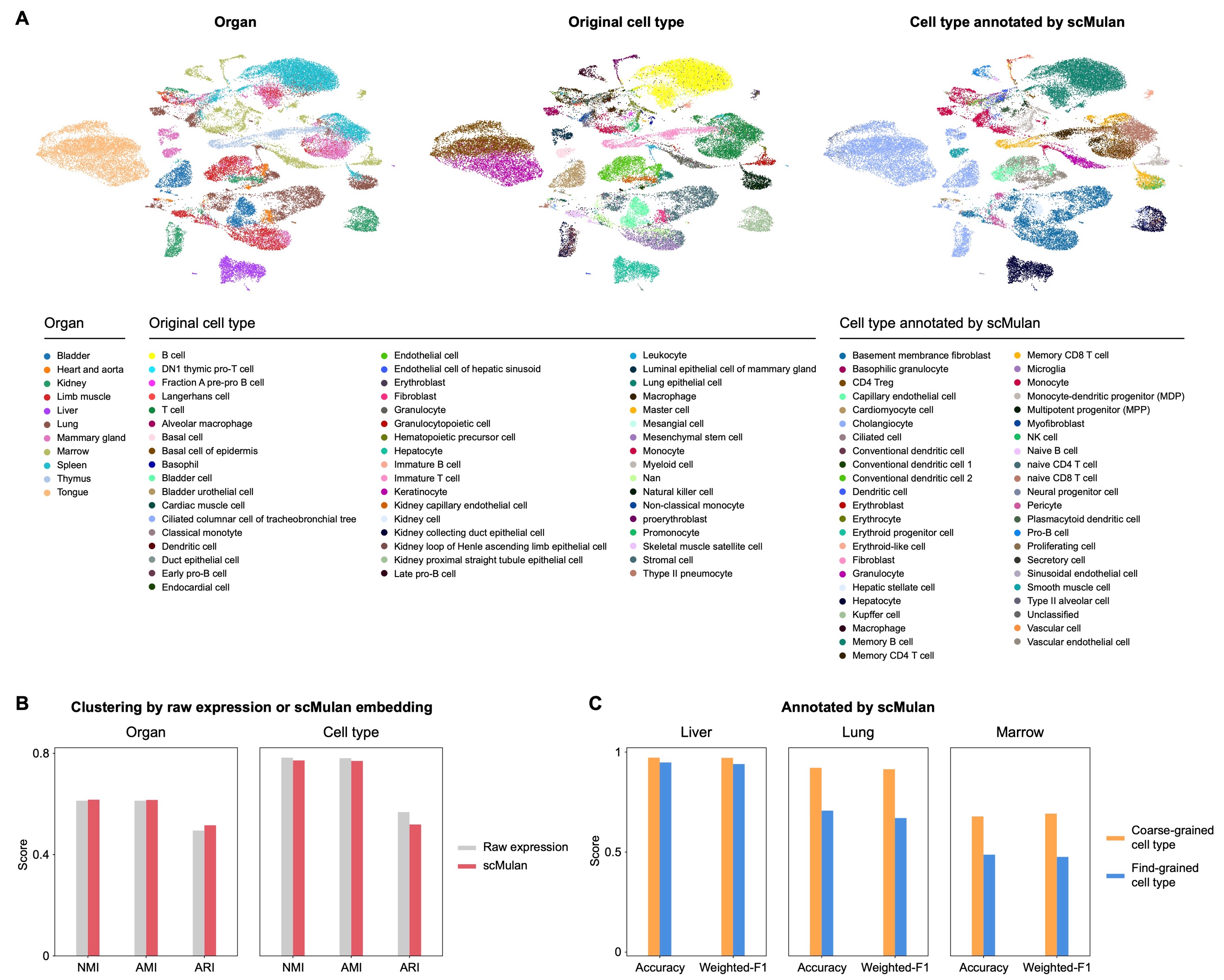

**Fig. S7. Performance of scMulan on annotating the Tabula Muris dataset.** (**A**) UMAP visualization of the scMulan embeddings. Cells are colored by organs, original cell types, and cell types annotated by scMulan. (**B**) Performance of scMulan embeddings on cell type clustering in comparison with raw gene expression profiles. (**C**) Performance of scMulan on cell type annotation.

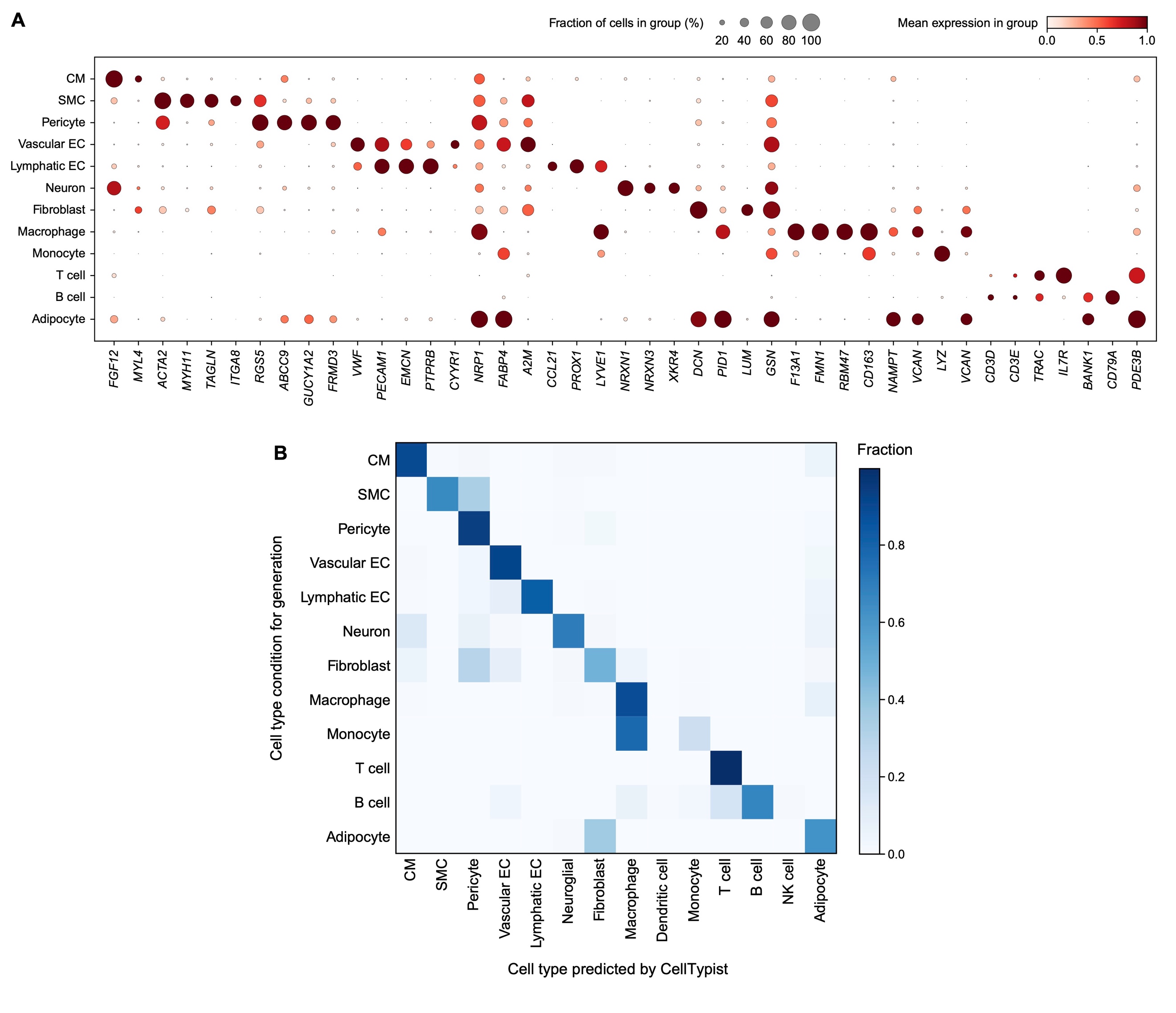

**Fig. S8. Expression patterns of the generated heart data.** (**A**) The expression profiles of marker genes on the generated heart data. (**B**) Automatic annotation results by CellTypist on the generated heart data. CM, cardiomyocyte cell/cardiac muscle cell; SMC, smooth muscle cell; EC, endothelial cell.

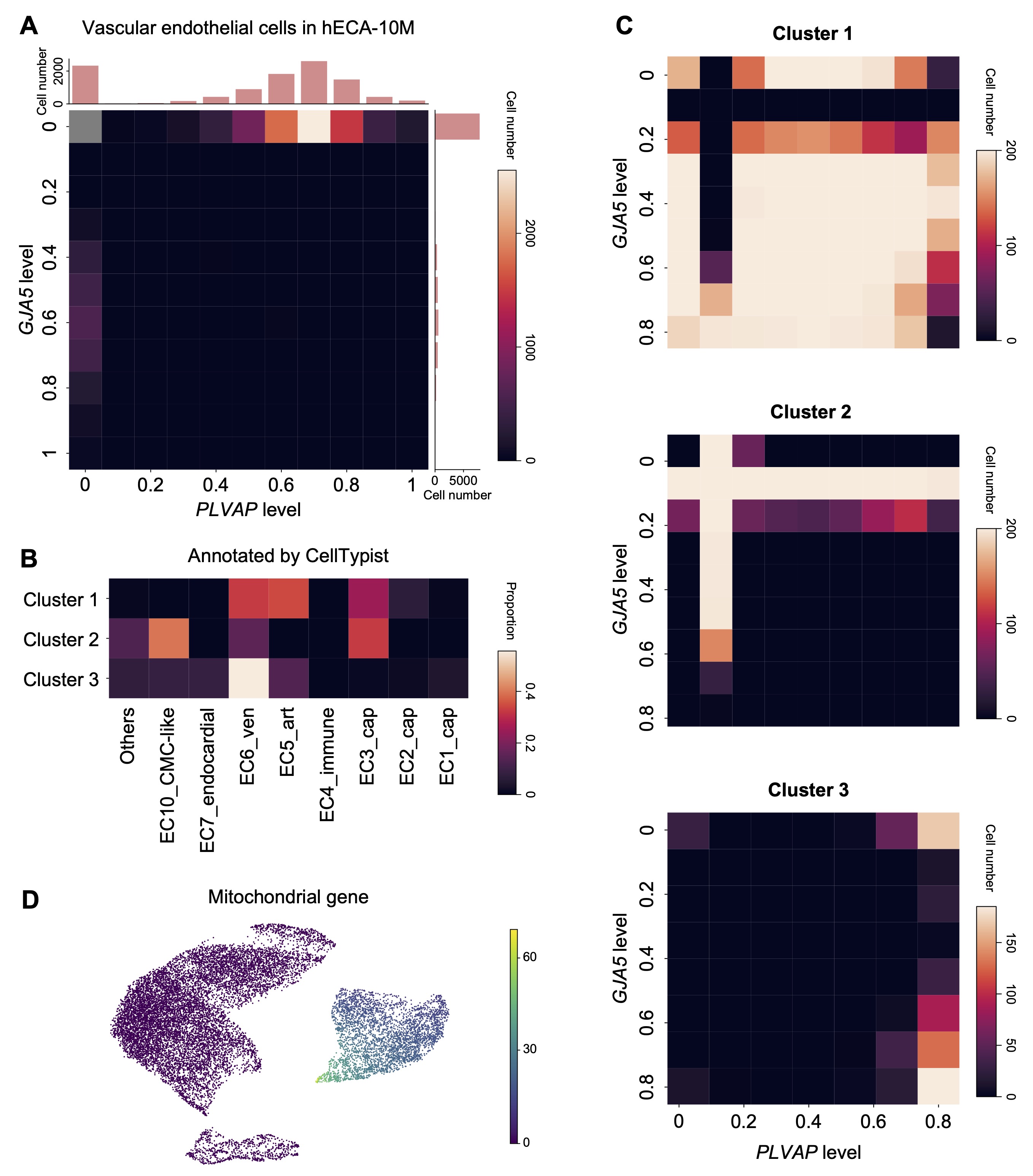

**Fig. S9. Expression patterns of the generated vascular endothelial cells.** (**A**) Expression patterns of *GJA5* and *PLVAP* of vascular endothelial cells in hECA-10M. (**B**) Automatic annotation results by CellTypist of the generated cells. (**C**) Expression patterns of *GJA5* and *PLVAP* of three clusters in the generated cells. (**D**) Mitochondrial gene expression in the generated cells.
